# Reticulate speciation and adaptive introgression in the *Anopheles gambiae* species complex

**DOI:** 10.1101/009837

**Authors:** Jacob E. Crawford, Michelle M. Riehle, Wamdaogo M. Guelbeogo, Awa Gneme, N’Fale Sagnon, Kenneth D. Vernick, Rasmus Nielsen, Brian P. Lazzaro

## Abstract

*Anopheles gambiae*, the primary vector of human malaria in sub-Saharan Africa, exists as a series of ecologically specialized subgroups that are phylogenetically nested within the *Anopheles gambiae* species complex. These species and subgroups exhibit varying degrees of reproductive isolation, sometimes recognized as distinct subspecies. We have sequenced 32 complete genomes from field-captured individuals of *Anopheles gambiae*, *Anopheles gambiae* M form (recently named *A. coluzzii*), sister species *A. arabiensis*, and the recently discovered “GOUNDRY” subgroup of *A*. *gambiae* that is highly susceptible to *Plasmodium*. Amidst a backdrop of strong reproductive isolation and adaptive differentiation, we find evidence for introgression of autosomal chromosomal regions among species and subgroups, some of which have facilitated adaptation. The X chromosome, however, is strongly differentiated among all species and subgroups, pointing to a disproportionately large effect of X chromosome genes in driving speciation among anophelines. Strikingly, we find that autosomal introgression has occurred from contemporary hybridization among *A. gambiae* and *A. arabiensis* despite strong divergence (∽5× higher than autosomal divergence) and isolation on the X chromosome. We find a large region of the X chromosome that has swept to fixation in the GOUNDRY subgroup within the last 100 years, which may be an inversion that serves as a partial barrier to contemporary gene flow. We show that speciation with gene flow results in genomic mosaicism of divergence and introgression. Such a reticulate gene pool connecting vector species and subgroups across the speciation continuum has important implications for malaria control efforts.

**Author Summary:** Subdivision of species into ecological specialized subgroups allows organisms to access a wider variety of environments and sometimes leads to the formation of species complexes. Adaptation to distinct environments tends to result in differentiation among closely related subgroups, although hybridization can facilitate sharing of globally adaptive alleles. Here, we show that differentiation and hybridization have acted in parallel in a species complex of *Anopheles* mosquitoes that vector human malaria. In particular, we show that extensive adaptive differentiation and partial reproductive isolation has led to genomic differentiation among mosquito species and subgroups, especially on the X chromosome. However, we also find evidence for exchange of genes on the autosomes that has provided the raw material for recent rapid adaptation. For example, we show that *A. arabiensis* has shared a mutation conferring insecticide resistance with two subgroups of *A. gambiae* within the last 60 years, illustrating the fluid nature of species boundaries among even more advanced species pairs. Our results underscore the expected challenges in deploying vector-based disease control strategies since many of the world’s most devastating human pathogens are transmitted by arthropod species complexes.

## Introduction

Closely related, morphologically similar, and sometimes interbreeding species complexes are common in nature and challenge conceptions of species boundaries [1]. In some cases, taxa diverge to exploit distinct ecological niches and thus represent incipient species on a trajectory towards reproductive isolation and phenotypic differentiation [2]. However, an alternative, more fluid, view of species boundaries and divergence as a continuum may be more appropriate in cases where the extent of reproductive isolation varies across time and environmental space (reviewed in [3]). Whether permeable species boundaries affect adaptive evolution among closely related taxa remains contentious. On one hand, introgression may serve to homogenize diverging taxa and oppose adaptive differentiation [4]. On the other hand, globally adaptive alleles may be shared among subgroups, increasing the mean fitness of each [5,6]. In species complexes, semi-permeable species boundaries could provide conduits for adaptive alleles to spread across large distances, both geographic and environmental.

The *Anopheles gambiae* species complex in sub-Saharan Africa includes several major vectors of malaria, which continues to place a devastating burden on local human populations [7]. Prior to the 1940s, *A. gambiae* was considered a single biologically variable species, but crossing studies and genetic analysis led to the naming of nine morphologically similar species that vary in their geographic distribution and ecology (reviewed in [8]). It is becoming increasingly appreciated from ecological distinctions and recent discoveries of additional genetic substructure that, even within species, *Anopheles* species frequently form partially reproductively isolated and differentiated subpopulations [9–11].

The M and S “molecular forms” are two major subgroups within *Anopheles gambiae sensu stricto* that are mostly reproductively isolated in the field, although they are compatible in captivity [12,13]. The two forms are sympatric in West and Central Africa but exploit distinct microecological niches [9,14]. The M form was recently given a formal species name, *Anopheles coluzzi* [8], although in the present paper we use the older terminology of ‘M form’ in the present work for continuity with existing literature. The ‘S form’ retains the name *A. gambiae s.s.*, but we use ‘S form’ here. Internal subdivisions exist even within the molecular forms [15,16] and recent evidence indicates an often high level of local M-S hybridization[11], illustrating the fluid semispecies boundaries within the *Anopheles gambiae* group. It is not known how many cryptic subpopulations exist within *Anopheles*, or how much gene flow they share, but there is evidence that subdivision may be common [11]. This is exemplified by the recently discovery of the GOUNDRY subgroup in Burkina Faso, which shows considerable genetic distinction from other described *A. gambiae* and is particularly permissive for *Plasmodium* development [17].

Epidemiological modeling and vector-based malaria control strategies must explicitly consider unique aspects of subgroups such as M, S, and GOUNDRY if they are to effectively predict disease dynamics and responses to intervention [18]. Because many of the world’s most devastating human diseases are vectored by arthropods in species complexes, including *Anopheles* vectors of malaria, tsetse fly vectors of African sleeping sickness, and *Ixodes* tick vectors of Lyme disease [19–21], the dynamic evolution of phenotypic diversity and adaptive introgression among cryptic taxa in species complexes has serious implications for public health. In the specific case of malaria, the existence of partially differentiated but occasionally hybridizing subspecies could complicate malaria control efforts that rely on the spread of transgenes through mosquito populations or conventional controls that target specific aspects of mosquito ecology or life history [22].

Genomic dissection of species complexes also lends insight into processes of speciation. Studies have shown strong differences between sex chromosomes (X or Z) and autosomes in the rates of genetic divergence and introgression and therefore speciation among pairs of species in systems ranging from *Drosophila* to Hominids. Genetic divergence tends to be higher on sex chromosomes relative to autosomes and introgressed regions are underrepresented on the sex chromosomes [23–25]. Heterogeneity in levels of genetic divergence has also been shown among genomic regions that vary in rates of meiotic recombination, with elevated divergence in genomic regions with low recombination rates such as centromeric regions and chromosomal inversions [26–28]. Analysis of genetic divergence and introgression in the *Anopheles gambiae* species complex provides a valuable opportunity to test the generality of these patterns across the speciation continuum.

The *Anopheles* system has become a model system for studying speciation and gene flow. Although substantial effort has been dedicated to understanding the status, history, and genomic consequences of reproductive isolation between two members of the *Anopheles gambiae* species complex, the M and S molecular forms of *A. gambiae* [13,29–31], firm conclusions regarding the degree of reproductive isolation and the age of the two forms remains controversial. This is due in part to the fact that these studies were based on ascertained SNP-panels, low-resolution sequencing datasets, and sequence datasets from small samples of laboratory mosquito colonies. In addition, they relied on statistical approaches that are incapable of distinguishing between multiple confounding population genetic processes. In particular, these analyses relied on measures of relative divergence, which are not robust to variation in recombination rates and natural selection across the genome and do not explicitly distinguish between lineage sorting of ancestral polymorphism and introgression [32–35]. Relatively few studies have addressed divergence among other members of the species complex and the same analytical concerns apply here as well [30,36,37]. In depth analysis of full sequence data using appropriate statistical approaches will greatly improve our ability to infer the introgression dynamics and the evolutionary consequences.

In the present study, we generated whole genome sequences from 32 wild caught females from the *Anopheles gambiae* species complex representing multiple points along the speciation continuum within this complex. We conducted detailed population genetic analysis of these data and show that 1) *A. gambiae* is more closely related to *A. arabiensis* than to *A. merus*, and the newly discovered GOUNDRY subgroup of *A. gambiae* diverged from the M form ∽100 kya, 2) speciation is driven by the X chromosome in this species complex, but introgression of autosomal regions is common among species across the speciation continuum, 3) ecological specialization and local adaptation may have driven many recent selective sweeps in these species, and 4) introgression has facilitated adaptation in some cases.

## Results

### Genome Sequencing and Population Genetic Analysis

We have completely sequenced the genomes of 32 field-captured female *Anopheles* mosquitoes from Burkina Faso and Guinea using the Illumina HiSeq2000 platform. We sequenced *A. gambiae* GOUNDRY (n=12), *A. gambiae* M form (n=10), *A. gambiae* S form (n=1) and *Anopheles arabiensis* (n=9). Most individuals were sequenced to an average read depth of 9.79x, while one individual each from GOUNDRY, M form, and S form was sequenced to at least 16.44x (Table S1). We also used publicly available genome sequences from *Anophleles merus* as an outgroup and a previously published *An. arabiensis* genome from Tanzania for a geographically distinct comparison [38]. We conducted population genetic analysis of aligned short-read data using genotype likelihoods and genotype calls calculated using the probabilistic inference framework ANGSD [39].

### Genetic Relatedness Among Species and Subgroups

To confirm broad-scale genetic relationships among the *Anopheles* included in this analysis, we calculated an unrooted neighbor-joining tree based on genome-wide genetic distance (*D*_*xy*_) at intergenic sites (Figure 1). Comparisons of genetic distance averaged across the genomes in our data recovers monophyletic species and subgroups with the *A. gambiae* clade more closely related to *A. arabiensis* than either is to *A. merus*, consistent with recent analyses [37,40]. Specifically, the length of the branch leading to the *A. merus* clade (0.0240 ± 7.61×10^−4^) is 3.64× the length of the branch leading to the *A. arabiensis* clade (0.0066 ± 7.41×10^−4^) and their 95% bootstrap intervals do not overlap indicating a closer genetic relationship between *A. arabiensis* and *A. gambiae*. In contrast to the initial description of the recently discovered GOUNDRY subgroup of *A. gambiae* as a genetic outgroup to the M and S molecular forms of *A. gambiae* [17], our data indicate that GOUNDRY is actually derived from the M form (*D*_*GM*_ = 0.0109; 100% bootstrap support) more recently than either subgroup shares a common ancestor with the S form (*D*_*GS*_ = 0.0149; *D*_*MS*_ = 0.0143).

**Figure 1:**
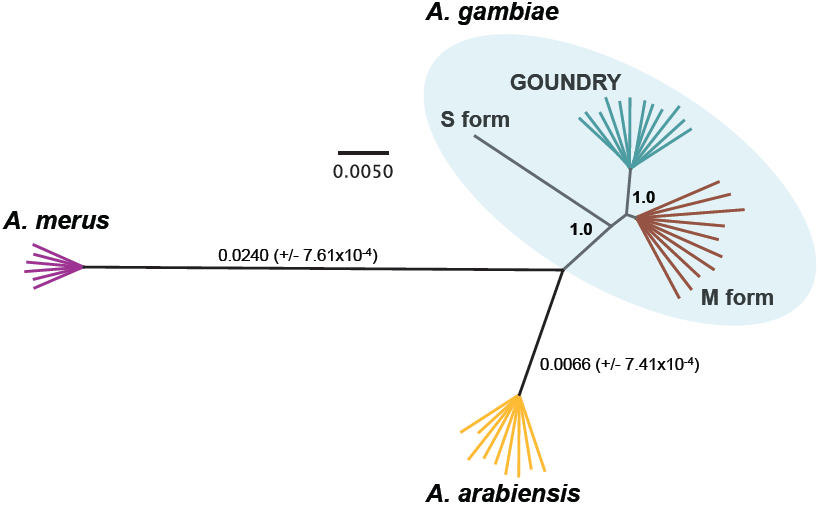
Average genetic relationships among species and subgroups in *Anopheles gambiae* species complex. Unrooted neighbor-joining tree calculated with the *ape* package in R and drawn with Geneious software. Branches indicate genetic distance (*D*_*xy*_) calculated using intergenic sites (see Methods) with scale bar for reference. Bootstrap support indicated for nodes leading to *A. gambiae* clade and GOUNDRY-M form clade. Branch lengths and 95%CIs indicated for branches leading to *A. merus* and *A. arabiensis*.

### Origins of GOUNDRY, a New Cryptic Taxon

It has been hypothesized that the advent of agriculture in sub-Saharan Africa ∽5-10 kya played a role in driving diversification and expansion of *Anopheles* mosquitoes [41]. To test whether the origin of GOUNDRY could have been associated with habitat modification driven by agriculture, we fit three-epoch population historical models (Figure 2; Methods) to the two-dimensional site frequency spectrum for GOUNDRY and M form *A. gambiae* using *dadi* [42]. The best-fitting model (Table 1) predicts that these subgroups diverged ∽111,200 ya (95% CI 96,718 – 125,010), followed by a 100-fold reduction in the size of both subgroups after isolation (Methods), and thus rejects any role of modern agriculture in subgroup division, although it should be noted that estimates of such old splits times inherently carry considerable uncertainty. Our inferred model is inconsistent by an order of magnitude with agriculture as a driving force in cladogenesis and more consistent with habitat fragmentation and loss due to natural causes. The model supports a >500-fold population growth in M form and 19-fold growth in GOUNDRY with extensive gene flow between them ∽85,300 ya, consistent with a re-establishment of contiguous habitat and abundant availability of bloodmeal hosts. Interestingly, the model supports additional population growth in both subgroups in the most recent epoch, which spans the last 10,000 years and coincides with the advent of agriculture. Hybridization related to secondary contact has not led to complete homogenization, however, as we conservatively identified nearly 9,000 fixed nucleotide differences distributed across the genomes of the two subgroups.

**Table 1:** Optimized parameter values and confidence intervals from best fit demographic model for GOUNDRY and M form. See Figure S2 for parameter descriptions.

| Parameter | Optimized Value | 95% Confidence Interval |  |
| --- | --- | --- | --- |
|  |  | Lower | Upper |
| $\theta (4N_A\mu L)^1$ | 180,914 | | |
| $N_A^2$ | 126,252 | 88,916 | 150,976 |
| ***Split Times*** |  |  |  |
| $t_1$ | 259,220 | 114,087 | 396,171 |
| $t_2$ | 754,204 | 741,946 | 766,215 |
| $t_3$ | 99,235 | 93,113 | 105,754 |
| $t_{TOT}$ | 1,112,660 | 967,181 | 1,250,106 |
| ***Population sizes*** |  |  |  |
| $N_{M1}/N_A$ | 0.01 | 0.01 | 0.01 |
| $N_{M2}/N_A$ | 5.74 | 4.58 | 7.65 |
| $N_{M3}/N_A$ | 12.34 | 9.54 | 16.61 |
| $N_{G1}/N_A$ | 0.01 | 0.00* | 0.08 |
| $N_{G2}/N_A$ | 0.19 | 0.11 | 0.32 |
| $N_{G3}/N_A$ | 0.78 | 0.62 | 1.05 |
| ***Migration rates***<br>***(4Nm)<sup>3</sup>*** |  |  |  |
| G1 into M1 | 0.010 | 0.0047 | 0.02 |
| G2 into M2 | 1.29 | 1.25 | 1.33 |
| G3 into M3 | 0.37 | 0.08 | 0.74 |
| M1 into G1 | 0.080 | 0.00* | 0.64 |
| M2 into G2 | $1.17 \times 10^{-4}$ | 0.00* | 0.0015 |
| M3 into G3 | 1.50 | 1.44 | 1.55 |
| 1 – Instead of estimating a confidence interval for $\theta$ which itself is not model parameter, we solved for $N_A$ and calculated a confidence for this parameter. In the implementation of *dadi*, $N_A$ is used to scale all other parameters in the model.<br>2 – Ancestral population size was calculated from the estimate of $\theta$ by dividing this value by 4 times the number of sites times the mutation rate (see Methods above).<br>3 – Values of $4Nm$ were calculated by multiplying the migration rate reported by *dadi* ($2N_{Am}$) by 2 times the ratio of the effective size of the recipient population (e.g. $N_{G1}$) over $N_A$.<br>* indicates cases where the lower bound of the 95% CI was negative. This is not meaningful, so we set these values to 0. |  |  |  |

**Figure 2:**
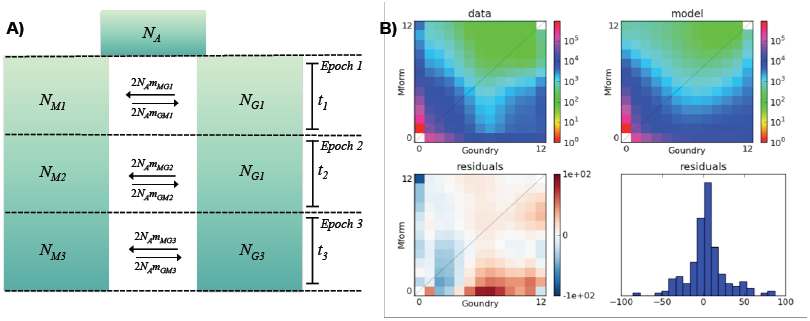
Demographic history of GOUNDRY and M form *A. gambiae*. **A)** Three-epoch demographic model. *N* parameters indicate effective population sizes. The duration of each epoch is indicated with the *t* parameters. Migration parameters (2*Nm*) are included as functions of the ancestral effective size. We included separate migration parameters for M into GOUNDRY migration (2*N*_*A*_*m*_GM_) and GOUNDRY into M (2*N*_*A*_*m*_GM_). **B)** Autosomal two-dimensional site-frequency spectra for GOUNDRY and M form for both the data and model. Residuals are calculated as the normalized difference between the model and the data (model – data), such that red colors indicate an excess number of SNPs predicted by the model.

We note that the dates reported here depend on assumptions about both the physiological mutation rate as well as the number of generations per year, neither of which are well known in *Anopheles* (Methods). As such, the details of these results would differ somewhat if different estimates were used, but we would have to invoke extreme values of these parameters in order to obtain estimates of an age of the GOUNDRY-M form split that is consistent with the advent of agriculture. The resulting 2D spectra from the data and the best-fit model are presented in Figure 2. Residuals indicating differences between the model and data are also presented. In general the fit is quite good. Most of the weight in the 2D empirical spectrum is matched in the 2D spectrum from the model, and the residual plot reveals that in general the model captures most features of the true spectrum. Several exceptions exist, however, indicating that some complexity of the true history is not captured by our model. Nonetheless, the model captures the patterns in the data quite well and suggest that GOUNDRY and M form diverged ∽100 kya, have both undergone bouts of population growth and experienced increased rates of hybridization in more recent evolutionary time.

### Evidence for extensive adaptive differentiation

Ecological specialization typically involves natural selection on ecologically relevant traits involving one or more genetic loci. To identify loci putatively involved in ecological specialization and quantify the role of selection in adaptive differentiation, we scanned the genomes of GOUNDRY, M form, and *A. arabiensis* for signals of recent positive natural selection using SweepFinder [43]. We evaluated the global site frequency spectra from intergenic sites in euchromatic regions of the autosome and X chromosome separately without explicitly calling genotypes using algorithms within ANGSD [39,44–46]. For M form and GOUNDRY, we established genome-wide critical values by comparing empirical likelihood ratio scores to a distribution of high scores obtained by applying SweepFinder to genomes simulated with coalescent simulations under the demographic model inferred for these subgroups described in the section above. For *A. arabiensis*, we used a ‘high-score-cluster’ approach to identify putative sweeps where regions were considered significant if two adjacent windows were assigned likelihood ratio scores from the top 0.1% genome-wide (see Methods). Consistent with ongoing adaptive divergence, perhaps related to ecological specialization, we find evidence for a number of recent selective sweeps that are private to each of the three groups [Figure 3, Tables S2 and S3]. We find evidence of 120 recent selective sweeps in the M form subgroup, compared to only 67 in GOUNDRY and 33 in the *A. arabiensis* genome.

**Figure 3:**
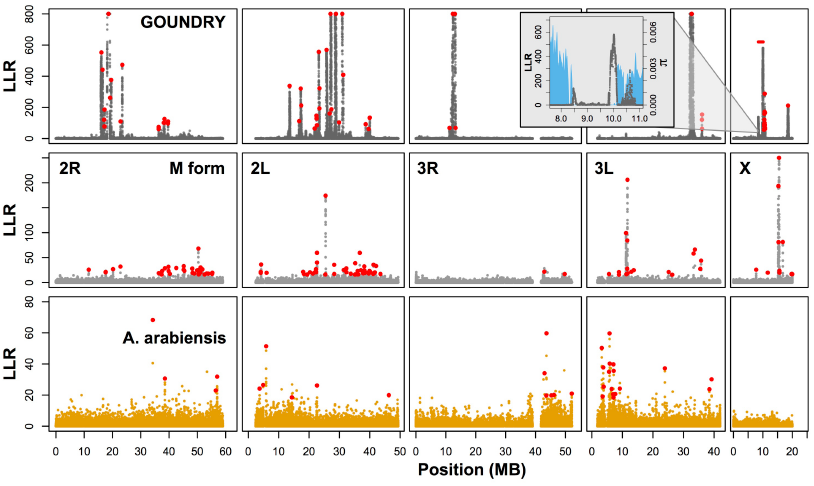
Maps of recent positive selection indicate adaptive differentiation in GOUNDRY (top row), M form (middle row), and *A. arabiensis* (bottom row). Recent selective sweeps are private and distributed across the genome of each group. Positive selection is significantly more prevalent in M form than GOUNDRY (*P* < 0.0001). Each point indicates the Log-Likelihood Ratio (LLR) value for a selective sweep at a given genomic position. Credible peaks are indicated with red dots. The large sweep region on the X chromosome in GOUNDRY is indicated with a horizontal red line, and the inset shows both LLRs and nucleotide diversity (blue). GOUNDRY windows with LLR values > 800 are truncated for presentation. Low-complexity regions were excluded. A full list of inferred targets of selection is given in Tables S2 and S3.

We were interested in determining whether GOUNDRY or M form has experienced significantly more selection than the other. We scanned each genome for selective windows and found that, of the 230 possible megabase windows, 36 windows harbored at least one selective sweep in GOUNDRY while 58 harbored sweeps in M form, indicating that M form has experienced significantly (Figure S1; *P* < 0.0001) more positive selection in recent evolutionary history. This difference may reflect recent exploitation of marginal ecological habitats by the M form subgroup, especially those associated with human settlements [9,47].

We note here that the many recent selective sweeps in GOUNDRY combined with the fact that GOUNDRY is partially inbred have led to unusual patterns of nucleotide diversity across the genome (Figure 4). We observed strong evidence of inbreeding in the GOUNDRY sample, but not in the other subgroups. Optimized individual inbreeding coefficients for chromosomal arms ranged from 0 to nearly 0.7 in the *A. gambiae* GOUNDRY subgroup sample, where an inbreeding coefficient of 1 would indicate complete lack of heterozygosity. Other than relatively high values on 3R for *A. arabiensis*, inbreeding coefficients for the M form subgroup and *A. arabiensis* were uniformly low. The underlying source of the high values on 3R in *A. arabiensis* is not known. In GOUNDRY, the homozygous regions are uniformly distributed around the genome but unique to each sampled individual, which is consistent with a high degree of close inbreeding. Importantly, inbreeding affects the distribution of diversity along diploid genomes, but should not obscure the signals of selective sweeps observed in GOUNDRY. Specifically, selective sweeps involve the fixation of the same haplotype across the population, whereas inbreeding simply brings together parental haplotypes that are identical-by-descent but are not necessarily shared among breeding pairs. As such, these processes have distinct effects on patterns of polymorphism that should not impact the ability to detect sweeps.

**Figure 4:**
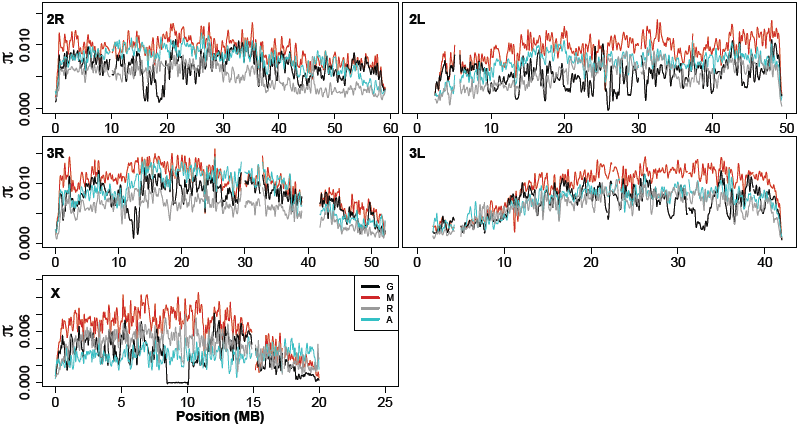
Chromosomal distributions of nucleotide diversity (π) at inter-genic sites (LOESS-smoothed with span of 1% using 10 kb non-overlapping windows). Low complexity and heterochromatic regions were excluded. M = *A. gambiae* M form; G = *A. gambiae* GOUNDRY; R = *A. merus*; A = *A. arabiensis.*

Two of the swept loci in the M form [*TEP1* and *Resistance to dieldrin* (*Rdl*)] have been identified previously using independent datasets and analytical approaches [48][13], providing an important validation of our analysis pipeline. These genes are involved in immunity and resistance to insecticide, respectively. We additionally find a particularly intriguing novel sweep at a gene encoding a neuropeptide F (NPF; AGAP004122). In *Drosophila*, NPFs have been shown to control larval behavior and feeding [49], so it is tempting to speculate that adaptive differentiation of this gene may underlie the previously described unique larval bottom-feeding and extended predator avoidance behaviors of the M form [50]. The most notable signal of positive selection in M form genomes is found near MB 15 on the X chromosome. Linkage disequilibrium among SNPs at least 1,000 bp apart is exceptionally elevated across a region approximately one megabase in size, suggesting that this region has been subjected to especially strong and recent positive selection (Figure S2). However, it is difficult to say whether or not this was a single event since our SweepFinder analysis indicates five independent peaks in the log-likelihood ratio surface in this region (Table S2), even after re-analyzing clusters of peaks that could represent single events falsely divided into a swarm of smaller peaks (Methods). The genes nearest to the five peaks include a gene encoding a cuticular protein in the RR-2 family, a gene encoding CDC-like kinase, and two genes with no known function (Table S2).

The sweeps in the *A. arabiensis* genomes include genes encoding two proteins involved in development, Eclosion hormone and Frizzled-2, and the gene encoding the sulfonylurea receptor. RNAi knockdown of the sulfonylurea receptor in *Drosophila melanogaster* significantly increases susceptibility to viral infections [51].

One particularly notable sweep in GOUNDRY lies near the gene encoding the master gustatory and odorant receptor *Agam/Orco* (*AgOr7*) that is a required coreceptor for all odorant binding proteins [52]. Expression of *AgOr7* is predominantly localized in female antennae and palps and fluctuates with circadian cycles, leading to the hypothesis that this gene controls temporal shifts in olfactory sensitivity [53]. Adaptation at *AgOr7* could have important impacts on anthropophily, sensitivity to DEET [54], and may underlie exophilic behavior in GOUNDRY [17].

In summary, we identified a large number of recent bouts of positive selection in GOUNDRY, M form, and *A. arabiensis*, including a number of interesting loci that potentially underlie important ecological and physical differences between these taxa. We also showed that M form has experienced more recent positive selection than GOUNDRY, which may reflect the more diverse environmental challenges that the M form has experienced as it exploits novel environments and niches.

### Recent autosomal introgression

Since recently diverged subgroups and species may still share polymorphism even if they are completely reproductively isolated in practice, we addressed the question of whether several members of the *Anopheles gambiae* species complex have introgressed historically or contemporaneously by formally differentiating between introgression and lineage sorting of ancestral polymorphism. To explicitly differentiate between these two population genetic processes, we used the ABBA-BABA test [55,56]. This test uses the *D* statistic to compare the distribution of alleles on the four taxon tree ((H1,H2),H3),O), where H1 and H2 are sister taxa and H3 and O are the outgroups. Under the null hypothesis of a perfect tree structure and no gene flow, the number of derived mutations that are shared only between the genomes of H2 and H3 (ABBA) is expected to equal the number of those that are shared only between H1 and H3 (BABA). *D* is then calculated as the standardized difference between the numbers of ABBAs and the number of BABAs with an expectation that *D* is zero under the null hypothesis [55]. Significant excess sharing of derived alleles between H3 and either H1 or H2 will result in a non-zero *D* and provides evidence of introgression. In our case, such tests are not appropriate because the alternative to no introgression is not only asymmetric introgression between an outgroup and one of the ingroups, but also potentially approximately equal amounts of introgression between the outgroup and each of the two ingroups. For that reason we use a modified test in which we instead consider the length distribution of fragments of shared ancestry. Since haplotypes are broken down by recombination over successive generations, the length distribution of shared haplotypes among populations is informative regarding the time since the most recent introgression event [57,58]. After *t* generations, the mean length of a shared haplotype is approximately (*r*(1 − *m*)(*t* −1)^−1^, where *r* is the recombination rate and *m* is the proportion of introgressed individuals in the population [58]. Since *t* will be smaller for introgressed haplotypes than for shared ancestral haplotypes, the mean length of introgressed haplotypes will be longer resulting in longer clusters of excess ABBAs or BABAs. We used patterns of linkage disequilibrium to approximate this expectation and deviations from it by calculating variance in the *D* statistic among ∽250 Kb blocks, Var[*D*_*BLOCK*_], to detect physical clusters of correlated genomic segments consistent with an excess of long shared haplotypes, and compared it to its null distribution generated by permuting fragments of size ∽2.5 Kb in order to manually break up possible correlations (see Methods). If the observed value of Var[*D*_*BLOCK*_] is significantly larger than that predicted from permuted fragments of size ∽2.5 Kb, which is >10x the average length of the decay of linkage disequilibrium, we conclude that the genomes harbor introgressed haplotypes.

We applied this test using several four-taxon trees and find evidence for recent introgression among species and subgroups of the *A. gambiae* species complex. We calculated the *D* statistic using a four-taxon tree with M form and GOUNDRY as the sister taxa (H1 and H2, respectively), *A. merus* as the outgroup (O), and either S form or *A. arabiensis* as H3. After comparing the Var[*D*_*BLOCK*_] in the empirical data to Var[*D*_*BLOCK*_] from 10^4^ randomly permutated genomes, we find significant evidence for introgression between the M form-GOUNDRY clade and S form [*P* < 0.0001, Figure 5]. Using M form and GOUNDRY as the ingroups again, but *A. arabiensis* as the test taxon (H3), we also find that the Var[*D*_*BLOCK*_] is significantly larger than all 10^4^ permuted genomes [*P* < 0.0001, Figure 5]. In fact, Var[*D*_*BLOCK*_] is larger by a factor of 3.7 in the S form test (0.0122) and a factor of 6.7 *A. arabiensis* test (0.0268) than the largest value of the 10^4^ permuted genomes (Figure 5). These results indicate that the observation of shared polymorphism between *A. gambiae* and *A. arabiensis* and between *A. gambiae* M and S forms can be attributed in part to introgression.

**Figure 5:**
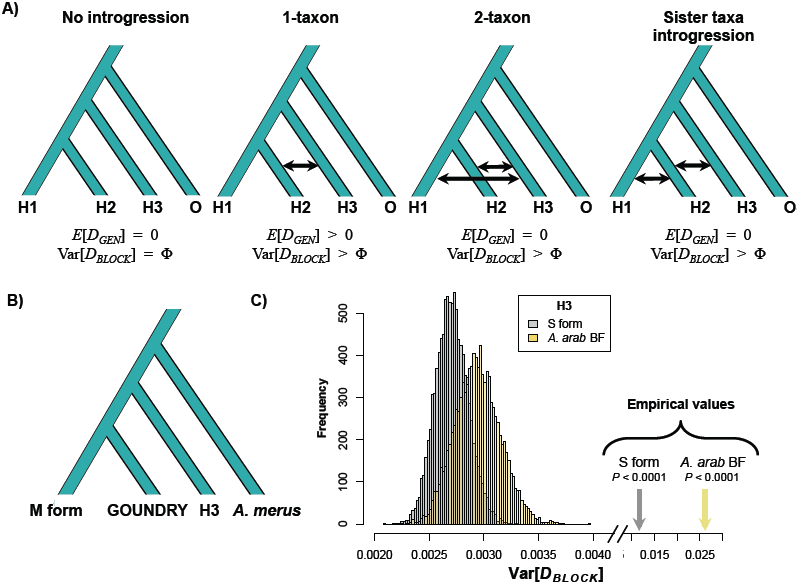
Excess variance among in *D*_*BLOCK*_ indicates recent introgression. **A)** Four-taxon trees used in ABBA-BABA tests with four alternative introgression models. The expected genome-wide value of the *D* statistic (*E*[*D*_*GEN*_]) is presented below in addition to the expected variance among *D* statistics calculated in genomic blocks (Var[*D*_*BLOCK*_]). Var[*D*_*BLOCK*_] under the ‘No introgression’ model is unknown and indicated here by *Φ* for comparison in other models. The ‘2-taxon’ and ‘Sister taxa introgression’ models may result in *E*[*D*_*GEN*_] of 0, but is expected to have increased Var[*D*_*BLOCK*_] relative to the ‘No introgression’ model providing a test for introgression. **B)** The four-taxon test tree used for analysis. **C)** The distributions of Var[*D*_*BLOCK*_] calculated from 10^4^ permuted genomes (see Methods) for test trees with *A. gambiae* S form (Grey) and *A. arabiensis* from Burkina Faso (Yellow) as the H3 outgroup are presented. The true Var[*D*_*BLOCK*_] values from each empirical data set, presented on a broken x-axis for comparison, are greater than all permuted genomes in each case consistent with the presence of introgressed haplotypes in these genomes.

### Contemporary introgression between A. gambiae and A. arabiensis

We can also ask whether introgression has occurred via contemporary hybridization by comparing signals of introgression with sympatric and allopatric populations. If introgression has occurred recently, we expect stronger affinity among sympatric relative to allopatric populations [33,59,60]. We tested whether introgression between *A. gambiae* M form and *A. arabiensis* has been recent with sympatric *A. arabiensis* versus allopatric *A. arabiensis* from Tanzania. In this case, we used the standard ABBA-BABA test since we are explicitly testing a simple ‘1-taxon’ model (Figure 5). We tested for introgression using a four-taxon tree of high-coverage individuals with the two *A. arabiensis* individuals as the ingroups (H1 and H2) with M form as the outgroup (H3) and find a significant excess of shared derived mutations between M form *A. gambiae* and sympatric *A. arabiensis* relative to allopatric *A. arabiensis* from Tanzania (*D* = -0.0542, Block Jackknife Z-score = -13.1533, *P* = 1.63×10^−39^; Table 2). Similarly, we used GOUNDRY as H3 and find evidence for significant excess introgression between sympatric *A. arabiensis* and GOUNDRY (*D* = - 0.0441, Block Jackknife Z-score = -11.7559, *P* = 6.58×10^−32^; Table 2). In line with recent evidence of contemporary hybridization between *A. gambiae* and *A. arabiensis* in Uganda [61], our results provide strong evidence that introgression continues to occur via contemporary hybridization in Burkina Faso as well that has the potential to impact the evolution of both ecologically and epidemiologically relevant traits.

**Table 2:** Modified block-based ABBA-BABA test of introgression. Genome-wide 95% thresholds and the proportions of the genome in significant windows are presented for three comparisons.

| H1<sup>a</sup> | H2<sup>a</sup> | H3<sup>a</sup> | Upper 95%<sup>b</sup> | Lower 95%<sup>b</sup> | Prop genome in sig M form Windows<sup>c</sup> | Prop genome in sig GOUNDRY Windows<sup>c</sup> |
| --- | --- | --- | --- | --- | --- | --- |
| M form | GOUNDRY | S form | 0.204 | -0.204 | 0.0114 | 0.0325 |
| M form | GOUNDRY | *A. arabiensis* (Burkina Faso) | 0.248 | -0.164 | 0.0364 | 0.0354 |
| M form | GOUNDRY | *A. arabiensis* (Tanzania) | 0.256 | -0.156 | 0.0323 | 0.0312 |

a – Taxonomic assignment in the ABBA-BABA test tree ((H1,H2),H3),O). b – Boundaries of the genome wide 95% threshold region estimated using block jackknife estimates of standard error within genomic regions after permutation (n = 10<sup>4</sup> replicates). c – For each comparison, windows exceeding the 95% thresholds were identified, and the sum of their length was compared to the total length of the *A. gambiae* PEST reference.

### Introgressed genes

Introgressed haplotypes may contain genes with potentially important phenotypic effects and variation along the genome in concentration of introgressed haplotypes can be indicative of barriers to introgression, so we partitioned the signal of introgression to identify introgressed chromosomal segments. To do so, we established genome-wide significance thresholds by comparing each empirical *D*_*BLOCK*_ value to the distribution of the most extreme *D*_*BLOCK*_ value from each of the 10^4^ permuted genomes (Methods). Since each *D*_*BLOCK*_ is polarized, we can identify windows that show significant introgression with each of the sister taxa (i.e. M form and GOUNDRY) and H3 (Figure 6). After conservatively correcting for multiple testing genome-wide, we find significant evidence of introgression between M form and both S form and *A. arabiensis*, and the proportions of the genome that are represented by significant windows are 1.1% and 3.6% with S form and *A. arabiensis,* respectively. These introgressed chromosomal blocks include 97 annotated protein coding sequences introgressed between the M and S forms and 543 introgressed between M form and *A. arabiensis* (Table S4). Moreover, we find strong evidence for introgression of the pericentromeric region on chromosome 2L between the M and S forms, which contrasts starkly with previous suggestions that this region may be a barrier to introgression and important for speciation between these taxa [13,29,30]. A recent study based on SNP-genotype data identified the same signal of introgression in this genomic region and attributed the high frequency of the introgressed haplotype to the sharing of an adaptive insecticide resistance allele at the *kdr* locus [62].

**Figure 6:**
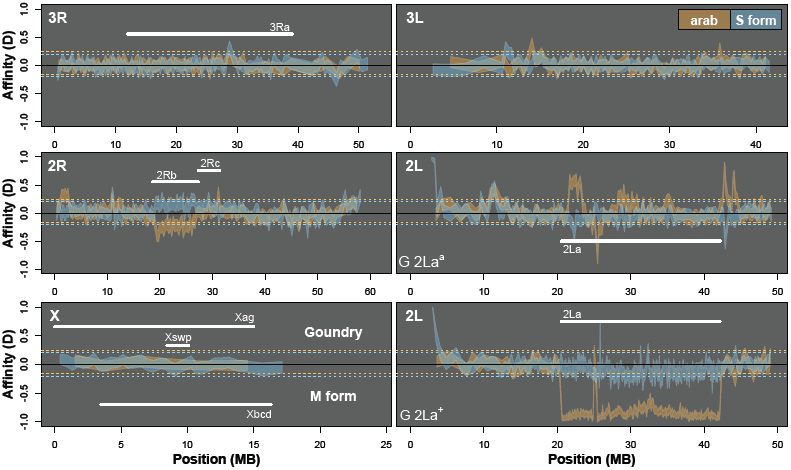
Significant autosomal introgression between *Anopheles* species and subspecies. ABBA-BABA statistics were calculated in non-overlapping windows of 500 informative sites using *A. merus* as the outgroup. Blue ribbon indicates 95% confidence region for introgression between S form (*2La^a/+^*) and GOUNDRY (positive *D*; *2La^a/a^* and *2La^+/+^*; *3R+; Xag*) and M form (negative *D*; *2La^a/a^*; *3R+*; *Xag*). Orange ribbon indicates 95% confidence region for introgression between *A. arabiensis* (*2La^a/a^*; *3Ra*; *Xbcd*) and GOUNDRY (positive *D*) and M form (negative *D*). Horizontal dotted lines (orange = *A. arabiensis*; blue = S form) indicate genome wide significance level after correction for multiple testing. Positions of relevant chromosomal inversions indicated with horizontal white lines. A full list of genes within significant windows is given in Table S4.

One window that shows an exceptionally strong introgression signal between M form and *A. arabiensis* harbors the GABA receptor gene, also known as the *resistance to dielrin* locus because of the role of this receptor in conferring resistance to the insecticide dieldrin and related insecticides [63]. Our finding that the *Rdl* locus has been introgressed between *A. gambiae* and *A. arabiensis* is contradictory to previous reports that these species have acquired resistance at this locus through independent but convergent mutations [64].

We used the same approach to identify genomic blocks shared between GOUNDRY and two H3 taxa, S form and *A. arabiensis* (Table 2). We find that windows representing 3.2% of the GOUNDRY genome and harboring 369 protein-coding sequences share a significant excess of derived mutations with the S form. In addition, we find that 3.5% of the GOUNDRY genome harboring 499 protein coding sequences shares a significant excess of derived mutations with *A. arabiensis*. In line with results above showing evidence of recent introgression among *A. gambiae* and sympatric *A. arabiensis* relative to allopatric *A. arabiensis*, we find that the windows with significant evidence of introgression with sympatric (Burkina Faso) *A. arabiensis* cover slightly more of the genome than windows with significant evidence of introgression with allopatric (Tanzania) *A. arabiensis* (Table 2). Although only relatively small percentages of these individual genomes show significant evidence of recent introgression, hundreds of protein-coding genes are involved in each case such that homogenization of these genomic regions may have large effects on phenotypic evolution, exhibited most prominently by the sharing of presumably functional mutations at two insecticide loci in the M form comparisons.

While identifying introgressed regions provides insight into the homogenizing effects of hybridization, identifying genomic regions depauperate of recent introgression can provide evidence for genomic barriers to introgression. Despite evidence for considerable introgression on the autosomes, we find no evidence of recent introgression of X chromosome sequence among any subgroups or species, which suggests a disproportionately large role for the X in speciation among these taxa.

### Introgressed chromosomal inversions

Partitioning the ABBA-BABA test into genomic blocks can also reveal possible evidence for introgressive sharing of chromosomal inversions (Figure 6). *A. arabiensis* is fixed for the inverted 2L*a*^*a*^ arrangement that spans ∽22 MB of chromosome 2L and is also nearly fixed in the subgroups of *A. gambiae* (i.e. the M and S forms) in most Savanna regions of West Africa [65]. However, both forms of the inversion (2L*a*^*a*^ and 2L*a*^+^) are segregating in GOUNDRY [17] and we analyzed both GOUNDRY forms here. Consistent with previous studies [30,65–68], we find strong evidence that the inverted form has introgressed between *A. gambiae* and *A. arabiensis.* The 2L*a* region exhibits extremely strong excess sharing of derived mutations when M form (2L*a*^*a*^) is H1, GOUNDRY (2L*a*^+^) is H2, and *A. arabiensis* is H3 (2L*a*^*a*^). This result reflects excess BABAs, or alleles shared between the 2L*a*^*a*^ haplotypes of M form and *A. arabiensis*. In a second comparison similar to the previous except that GOUNDRY is represented by its 2L*a*^*a*^ haplotype, the strong and extended reduction in *D* across the region is not observed. The strong reduction in *D* when the two inversion forms are contrasted provides evidence consistent with the introgression of the 2L*a*^*a*^ form of this inversion between *A. gambiae* and *A. arabiensis* before the split of M form and GOUNDRY. We also find a similar extended reduction in *D* on 2R coinciding with the known location of the 2R*b* chromosomal inversion (Figure 6). This reduction is also consistent with introgression of one form of this inversion between M form and *A. arabiensis,* although we don’t know the inversion karyotypes for the individuals analyzed here, so it is more difficult to interpret this signal. In principle, these results are consistent with introgression, but chromosomal inversions are known to suppress recombination to some extent [69–71] such that ancestral haplotypes may be maintained over time violating some of the assumptions our modified ABBA-BABA test. However, this confounding scenario can be ruled out in the case of 2L*a* since we find a strong reduction in genetic divergence (*D*_*xy*_) between the 2L*a*^*a*^ haplotypes of *A. arabiensis* and both M form and GOUNDRY relative to surrounding genomic regions (Figure 7), which is consistent with introgression but not ancestral haplotype maintenance. As a result, this region harbors genomic patterns of divergence and allele sharing that deviate from the genome at large in a comparison-specific fashion. Interestingly, we also find local deviations from this pattern that provide evidence for recent introgression events that are independent from the inversion introgression event (Figure 6). The combination of ABBA-BABA tests and genetic divergence among the chromosomal forms of these taxa provide strong evidence that 2L*a* has introgressed and suggestive evidence that 2R*b* may have crossed species boundaries.

**Figure 7:**
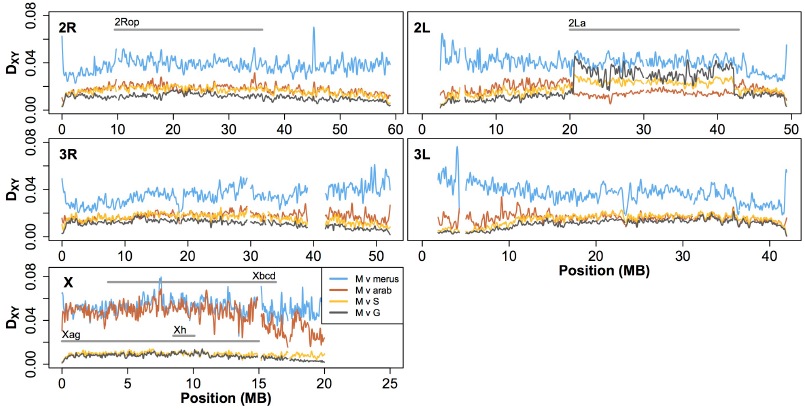
Patterns of divergence among subgroups of *A. gambiae* follow similar curves (LOESS-smoothed with span of 1% using 10 kb non-overlapping windows), although differing slightly in magnitude, except increases in pericentromeric region in the M vs. S comparison and inside the 2L*a* inversion where these populations differ in karyotype (G-2L*a*^+/+^, M-2L*a*^*a/a*^, S-2L*a*^*a/+*^). Divergence between the M form and *A. arabiensis* and *A. merus* is enriched on the X chromosome, especially inside the inverted X*ag* and *Xbcd* region (M vs. *arab*) and in pericentromeric regions (M vs. *merus*). Grey bars indicate locations of differentially fixed chromosomal inversions as well as the 2L*a* inversion and the large sweep on the GOUNDRY X (X*h*). Low complexity and heterochromatic regions were excluded.

### Introgression facilitates adaptation

Natural selection is expected to remove most introgressed genetic material due to ecological misfit or Dobzhanski-Muller Incompatibilities [72,73], especially between more distant species, but some introgressed alleles may be selectively favored in the recipient subgroup. As just described above, we find clear evidence for introgression of the large 2L*a* inversion between *A. gambiae* and *A. arabiensis*, which is in line with several previous studies [30,65,67,68]. A number of studies have associated this inversion with various putatively adaptive phenotypes and shown that the frequencies of 2L*a* karyotypes correlate strongly with environmental factors such as aridity in West and Central Africa [65,74]. Taken together, these observations indicate that the 2L*a*^*a*^ haplotype has crossed species boundaries and may have facilitated ecological and physiological adaptation.

To identify other putatively adaptive introgression events, we examined the intersection between our scans for selection and for introgression (both described above). Since we cannot determine the direction of introgression robustly with the framework used here, we identified selective sweeps in either the donor or recipient species and found 17 selective events that may be due to selection on alleles that were originally introgressed from another taxa. We find a large region on chromosome 2L (∽21.5-23.5 MB) that is enriched for introgression of derived mutations between GOUNDRY (2L*a*^*a*^) and *A. arabiensis* that harbors a recent sweep in *A. arabiensis* at the gene AGAP005855, which has no known function.

The most striking example of these putative adaptive introgression events involves allele sharing at the GABA receptor on 2L known to be involved in insecticide resistance in both *A. gambiae* and *A. arabiensis* ([64]; Figure 8). Our scan for selection points to a selective sweep in the M form subgroup at the boundary of the coding sequence for this gene (*P*_*gen*_ < 0.005, *P*_*loc*_ < 1.26×10^−06^). However, we also find evidence, albeit below the significance threshold at the genome level, for selection in both GOUNDRY and *A. arabiensis* (Figure 8). There is an independent signal of selection in GOUNDRY (*P*_*gen*_ > 0.05, *P*_*loc*_ < 2.67×10^−04^) upstream of the M form signal. We do not find LLR values above background levels that would be consistent with recent selection in *A. arabiensis*, but we find that LD is slightly elevated and nucleotide diversity reduced relative to neighboring regions, consistent with a historical sweep at this locus (Figure 8). It is important to note that the model implemented in SweepFinder is specifically aimed at detecting very recent and complete sweeps [43], so it is not surprising that we might see remnants of a putative historical sweep that is not picked up by SweepFinder. Together, these results suggest that a resistance allele arose first and adaptively fixed in *A. arabiensis*, and adaptive fixations occurred more recently in M form and GOUNDRY.

**Figure 8:**
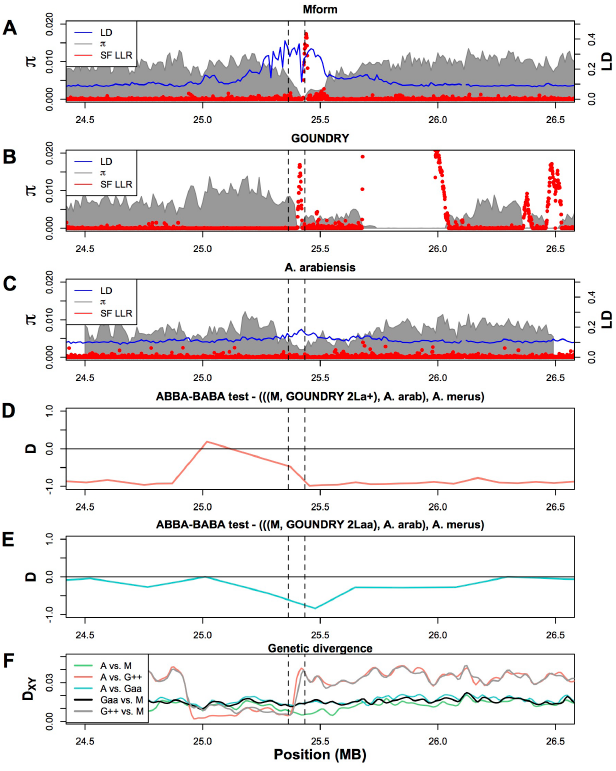
Patterns of nucleotide diversity (π, pi), background linkage disequilibrium (*r*^*2*^ SNPs > 1 kb apart; LD), genetic divergence (*D*_*xy*_), introgression (*D*), and selective sweeps (LLR) at the *Rdl* locus in *A. gambiae* M form, *A. gambiae* GOUNDRY, and *A. arabiensis*. Nucleotide diversity, LD, and SweepFinder LLRs are presented together for **A**) M form, **B**) GOUNDRY, and **C**) *A. arabiensis.* Y-axes are omitted for the SF LLR statistic. Linkage disequilibrium is not included for GOUNDRY because it could not be estimated robustly for GOUNDRY (see Methods). ABBA-BABA test statistics presented for the trees **D)** (((M form 2L*a*^*a*^, GOUNDRY 2L*a*^+^), *A. arabiensis* 2L*a*^*a*^), *A. merus*) and **E**) (((M form 2L*a*^*a*^,GOUNDRY 2L*a*^*a*^), *A. arabiensis* 2L*a*^*a*^), *A. merus*). **F)** Pairwise estimates of genetic divergence among *A. arabiensis* (A), M form (M), GOUNDRY 2L*a^a/a^* (Gaa), and GOUNDRY 2L*a^+/+^* (G++). Outer boundaries of the *Rdl* coding sequence are indicated with vertical dashed lines.

We find strong evidence that haplotypes involving the *Rdl* locus have been introgressed among *Anopheles.* It is necessary to consider the genomic background when interpreting data at the *Rdl* locus since if falls within the 2L*a* chromosomal inversion. On one hand genetic divergence is relatively low across the inverted 2L*a*^*a*^ region between *A. gambiae* and *A. arabiensis*, since this inversion has introgressed between these species. Conversely, there is exceptionally high genetic divergence between GOUNDRY 2L*a^+^* chromosomes and the M form subgroup (fixed for 2L*a*^*a*^ in this sample set) and *A. arabiensis* (fixed for 2L*a*^*a*^ as a species) chromosomes (Figure 7). Close inspection of ABBA-BABA tests and patterns of pairwise genetic divergence at the *Rdl* region suggests that both GOUNDRY and M form have independently received introgressed material in this region from *A. arabiensis* in a *2*L*a*-inversion karyotype dependent fashion. Specifically, the ABBA-BABA test involving the tree ((M form 2L*a*^*a*^, GOUNDRY 2L*a*^*a*^), *A. arabiensis* 2L*a*^*a*^), *A. merus* 2L*a*^*a*^) indicates evidence for significant allele sharing between *A. arabiensis* and M form at this locus (Figures 6 and 8). Together with the fact that genetic divergence between these two taxa is low at *Rdl* relative to immediately surrounding regions (Figure 8), these results provide strong evidence for introgression between M form and *A. arabiensis* at the *Rdl* locus. However, we also see evidence for an independent introgression event between *A. arabiensis* and the population of 2L*a^+^* chromosomes in GOUNDRY. In the ABBA-BABA test based on the tree ((M form 2L*a*^*a*^, GOUNDRY 2L*a^+^*), *A. arabiensis* 2L*a*^*a*^), *A. merus* 2L*a*^*a*^), the region inside the *2*L*a* inversion is almost entirely negative and nearly -1, reflecting the strong excess of allele sharing between the introgressed 2L*a*^*a*^ chromosomes from *A. arabiensis* and M form. However, amidst this pattern, we see a large region of positive *D* statistics at the *Rdl* locus indicating excess derived mutation sharing between *A. arabiensis* and GOUNDRY 2L*a^+^* chromosomes (Figures 6 and 8). Consistent with this result, we also see a large and extended reduction in genetic divergence between GOUNDRY 2L*a^+^* and both M form and *A. arabiensis*, indicating recent introgression between GOUNDRY and one of these groups. However, genetic divergence is exceptionally low between GOUDNRY and *A. arabiensis* across the entire putatively introgressed region, while divergence is reduced between M form and GOUNDRY only where genetic divergence is also low between M form and *A. arabiensis* (Figure 8). These observations provide evidence that an introgression event occurred first from *A. arabiensis* into M form followed by a second independent introgression event from *A. arabiensis* into GOUNDRY.

In summary, the evidence for positive selection at this locus in these groups in combination with the signals of independent introgression events lead us to hypothesize that a mutation conferring insecticide resistance arose and adaptively fixed first in *A. arabiensis* and then was shared successively with the M form and GOUNDRY subgroups where it underwent positive selection in these groups as well. Since insecticide pressure on mosquito populations in West Africa, and thus selective pressure on this variant, began in the 1950s [75], we infer that introgression of this adaptive allele among these taxa has occurred within the last 60 years.

### Xag-Xbcd inversion complex is barrier to introgression

We have shown above that some genomic regions have recently introgressed between the genomes of members of the *Anopheles gambiae* species complex, which raises the question of whether certain genomic regions contain barriers to such introgression. We hypothesized that long term differences in introgression along the genome will be reflected in patterns of genetic divergence, since genetic divergence will be partially determined in part by selection for and against introgressed material. Empirical and theoretical work has led to the hypothesis that hybrid sterility factors or locally adapted genetic variants in genomic regions with restricted recombination in hybrids play an especially important role in formation and maintenance of species boundaries in the face of gene flow because the hitchhiking process involved in selection against maladaptive introgressed variants will affect larger genomic swaths in lowly recombining regions [71,76–79]. In particular, chromosomal inversions and centromeric regions are thought to play a role in species formation when hybridization is ongoing, although currently available empirical evidence supporting a role of centromeric regions is confounded by the effects of natural selection on linked sites [32,33,35]. Empirical and theoretical evidence also supports the hypothesis that the X chromosome may play an important role in maintaining species boundaries (‘large-X effect’; [23,25,80,81]). To test for genomic barriers among taxa in the *Anopheles gambiae* species complex, we asked whether sequence divergence (*D*_*xy*_) between taxa at various distances along the speciation continuum differed among genomic regions as expected under a model of differential rates of introgression. However, *D*_*xy*_ is an estimator of 2*μt* + 4*Nμ*, where *μ* is the mutation rate, *t* is the number of generations since the species split, and *N* is the effective population size of the ancestral population, but 4*Nμ* can vary among genomic regions for reasons unrelated to differential introgression (i.e. variable mutation rate or effects of natural selection on linked sites; Figure 4). Therefore, we jointly analyzed nucleotide diversity (*π*) as a proxy for 4*Nμ* among genomic regions to avoid confounding intra-population effects with differential introgression (see Methods).

Hybrid male sterility maps to the X chromosome in *A. gambiae*-*A. arabiensis* crosses and two large X-linked chromosomal inversion complexes [X*ag* and X*bcd*] suppress recombination on the X in hybrids of these species, so we hypothesized that less frequent introgression may lead to exceptionally high sequence divergence on the X relative to other genomic regions. In support of this hypothesis, we found that nucleotide diversity on the X is significantly lower than on the autosomes (Mann-Whitney (hereafter M-W) *P* < 2.2×10^−16^), but that genetic divergence is significantly higher on the X (M-W *P* < 2.2×10^−16^; Figure 9). Moreover, genetic divergence is significantly higher in the region harboring inversions (M-W *P* < 2.2×10^−16^) than in the surrounding chromosome. Nucleotide diversity is also higher in this region relative to the centromere-proximal region (M-W *P* < 2.2×10^−16^), where nucleotide diversity is especially low, presumably due to the effects of linked selection on neutral genetic variation. While we cannot formally rule out an elevated mutation rate inside the inverted region, there is no reason to expect the inverted region to be more mutable. This excess genetic divergence on the X is consistent with our results in the ABBA-BABA analysis above showing that the X chromosome lacks evidence of recent introgression among these taxa. In general, these observations are in line with previous analyses of genetic differentiation among *A. arabiensis* and *A. gambiae*, as well as with laboratory backcrossing experiments [30,37,82], indicating that introgression is particularly inhibited on the X chromosome, and that the X chromosome plays a disproportionately large role in driving speciation.

**Figure 9:**
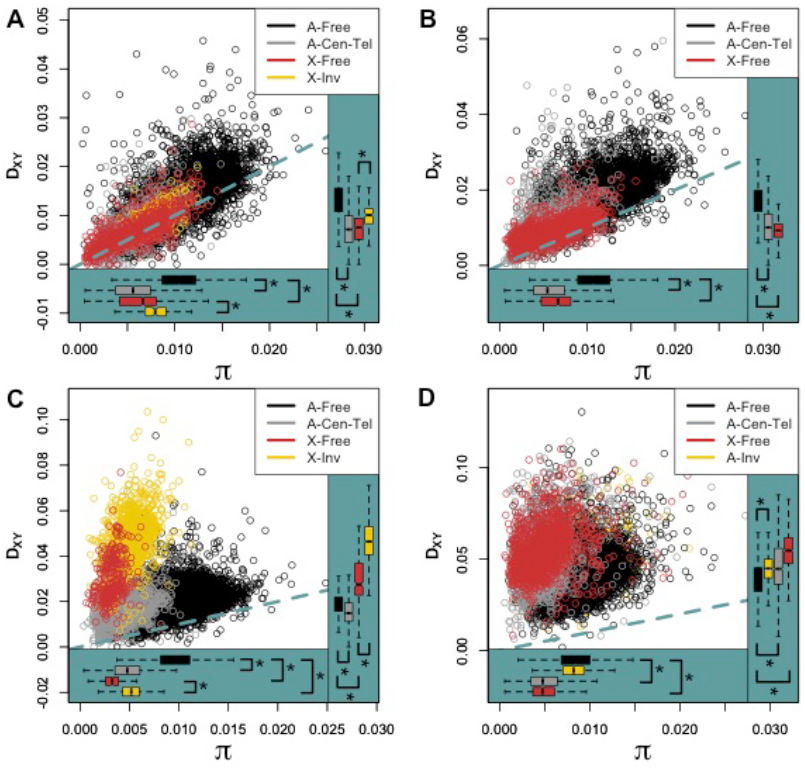
Patterns of genetic divergence (*D*_*xy*_) between populations as a function of nucleotide diversity (*π*) reveal differential gene flow during speciation. Genomic regions defined by expected rates of recombination in hybrids (see Methods) differ in their distributions of nucleotide diversity and genetic divergence, but not always in the same direction, suggesting that gene flow has been restricted on the X and lowly recombining regions in some cases. **A)** M form vs. GOUNDRY, **B)** M form vs. S form, **C)** M form vs. *A. arabiensis* **D)** M form vs. *A. merus*. Panel legends indicate colors corresponding to genomic location of each 10 Kb window where ‘Free’ indicates freely recombining regions, ‘Cen-Tel’ indicates centromeric and telomeric autosomal regions, and ‘Inv’ indicates chromosomal inversions. ‘A-’ and ‘X-’ indicate autosomal or X chromosome. Dashed blue-green line indicates perfect correlation. Asterisks indicate Mann-Whitney tests with *P* values < 3.92×10^−05^ for comparisons indicated with brackets. Note that the y-axis scale differs among panels.

In the comparison between M form and *A. arabiensis,* we generally find no evidence for elevated divergence across the autosome, indicating a long history of introgression and few genomic barriers. However, inspection of the genomic distribution of nucleotide diversity and divergence (Figures 4 and 7, respectively) reveals that the difference between nucleotide divergence and inter-species diversity is especially high in the pericentromeric region of chromosome 3. Nucleotide diversity is reduced in this region, presumably resulting in part from the effects of intra-population positive and negative selection on linked sites. However, the relative increase in *D*_*xy*_, which is not affected by positive selection, suggests that migrant alleles have been selected against in this region such that this region harbors both fewer shared polymorphisms as well as more private mutations relative to the nearby freely recombining regions, for example. This raises the question of what genetic factors might be involved in local adaptation or genetic incompatibilities in this region. We find evidence for adaptive divergence at multiple loci in this region, including one that involves the gene encoding *Frizzled-2*, a member of the *Wnt* receptor signaling pathway involved in development [83] that could underlie developmental divergence among these taxa.

Overall, we show that rates of introgression among *A. gambiae* and *A. arabiensis* have differed dramatically among genomic regions resulting in exceptionally high divergence on the X chromosome, intermediate divergence in the pericentromeric region of 3L, and divergence levels similar to subgroup comparisons within *A. gambiae* across most of the autosome. These data are consistent with the X chromosome playing a primary and disproportionately large role in speciation among these taxa and a secondary role for centromeric regions.

### Novel X-linked chromosomal inversion in GOUNDRY is a barrier to introgression

The most striking sweep from the selection scan of GOUNDRY genomes above covers 1.67 Mb on the X chromosome and results in nearly complete absence of polymorphism across this region (Figure 4), despite read depths in the region comparable to neighboring genomic regions (Figure S3). The remarkably large size of the region devoid of diversity would imply exceptionally strong positive selection under standard rates of meiotic recombination. For comparison, previously identified strong sweeps associated with insecticide resistance span approximately 40 Kb and 100 Kb in freely recombining genomic regions of *Drosophila melanogaster* and *D. simulans,* respectively [84,85]. The swept region in GOUNDRY is marked by especially sharp edges relative to the other footprints of selection in our data (Figures 3 and 4), implying that recombination has been suppressed at the boundaries this region. Collectively, these observations suggest that the swept region may contain a small chromosomal inversion, which we have named X*h* in keeping with inversion naming conventions in the *Anopheles* system. Notably, this pattern is virtually identical to the pattern of diversity in a confirmed X-linked inversion discovered in African populations of *D. melanogaster* [86]. The region in GOUNDRY includes 92 predicted protein coding sequences (Table S3), including the *white* gene, two members of the gene family encoding the TWDL cuticular protein family (*TWDL8* and *TWDL9*), and five genes annotated with immune function (*CLIPC4, CLIPC5, CLIPC6, CLIPC10, PGRPS1*). The lack of diversity in the region implies that the presumed X*h* inversion has a single recent origin and was quickly swept to fixation in GOUNDRY. We estimated the age of the haplotype (see Methods) inside the sweep region to be 78 years with a standard deviation of 9.15 by assuming that all mutations in the region are new and estimating the mean time since the most recent common ancestor of the sampled haplotypes, representing the time of the sweep. Such extraordinarily recent adaptation in an otherwise old subgroup is consistent with the selection pressures related to 19^th^ and 20^th^ century human activity such as insecticide pressure or widespread habitat modification.

We hypothesized that this putative X-linked chromosomal inversion in GOUNDRY may serve as a barrier to introgression with M form, which we showed in the demographic inference above has occurred at least in autosomal regions. We used the same logic as for the comparison with *A. arabiensis* to compare divergence among genomic regions and find that genetic divergence among M form and GOUNDRY generally scales with (M form) nucleotide diversity across the genome (Figure 9), consistent with broadly neutral divergence, suggesting that the rate of introgression among these subgroups is qualitatively similar genome-wide. Although the distributions of genetic divergence in 10 Kb windows are significantly different among genomic regions (M-W *P* < 2.2×10^−16^), the rank order of genetic divergence among genomic regions is the same as for nucleotide diversity in these regions. This suggests that differences in genetic divergence among regions can in large part be explained by differences in nucleotide diversity. However, the difference between the X chromosome at large and the putative X*h* inversion is slightly larger for the divergence measure than for nucleotide diversity, suggesting that divergence in the putative X inversion may be present that is too small to be detectable with this particular test of distributions.

Since we expect that the putative X*h* inversion is likely too young for measurable differences in the distributions of *D*_*xy*_ to have accumulated, we tested for excess divergence in the X*h* inversion using several additional more sensitive approaches. When divergence along the X chromosome is explicitly scaled by the mutation rate inferred from levels of polymorphism in the M form sample (*D*_*a*_), the putatively adaptive X*h* inversion between GOUNDRY and M form is proportionally much more divergent than is the remainder of the X chromosome (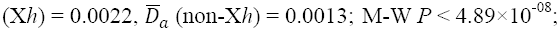; Figure S4). Although relative measures of divergence, such as *D*_*a*_, are known [32,33] to be confounded by reductions in nucleotide related to natural selection on linked sites, for example, we believe that this analysis is robust to these concerns because the comparison is among only X-linked windows and the region of interest is in a region of the chromosome that is highly diverse in subgroups where it has not been swept (Figure 4). We also directly tested whether the X*h* region is significantly more diverged than similarly sized windows on the X by defining a new statistic, *ΔD_xy_*, to compare the average *D*_*xy*_ among 10 kb windows within the putative inverted region (n = 166) with the average *D*_*xy*_ among windows in every other 1.67 MB window on the X that does not overlap with the inverted region (n = 1,499; Methods). A positive *ΔD_xy_* indicates that the average value inside the inversion is greater than the average value in the window outside the inversion. A negative value indicates that the average value is higher in the window outside the inversion. We calculated this statistic between M and GOUNDRY (see Methods) and found that *ΔD_xy_* was significantly positive (*P <* 10^−5^) for every window in the comparisons between M and GOUNDRY (Figure S5). Interestingly, the windows immediately adjacent to the inversion show the smallest *ΔD_xy_* values (Figure S5), suggesting that the introgression-limiting effects of the inversion have spread beyond the breakpoints. Both of these tests indicate that sequence divergence between M and GOUNDRY is greater inside the putative inversion relative to the X as a whole, which is not likely to reflect accumulation of many new private mutations inside the inversion, but rather a greater proportion of shared polymorphisms outside the inversion consistent with higher rates of introgression outside the inversion.

The effects of the putative X*h* inversion on the GOUNDRY X chromosome as a barrier to gene flow may extend beyond the boundaries of the inverted region, resulting in very recent accumulation of divergence on the X chromosome that is confounded in our analysis with inherent evolutionary differences between the autosomes and the X chromosome. One approach to estimate differences in divergence is to compare divergence between a focal pair of subgroups (GOUNDRY and M form) to divergence between one of the focal groups and an outgroup (GOUNDRY and S form) in order to scale by differences in mutation rate and other population genetic parameters among regions. This approach estimates what is known as Relative Node Depth (RND = *D*_*GM*_/*D*_*GS*_, where subscripts G, M, and S indicate GOUNDRY, M form, and S form respectively), and a higher RND indicates greater divergence between the focal groups [87]. We find that RND is 0.7797 on the autosomes and 0.8058 on the X, indicating lower relative genetic divergence between GOUNDRY and M form on the autosomes than on the X. To explicitly test whether such a pattern could be obtained under a pure split model with no gene flow, we obtained expected values of Relative Node Depth (RND) by assuming a phylogeny where M form and GOUNDRY form a clade with S form as the outgroup and using coalescent theory under this model to calculate expected RND (Methods).

Our analytical results support the hypothesis that *D*_*GM*_ is downwardly biased relative to *D*_*GS*_ on the autosomes as a result of higher rates of gene flow on the autosomes relative to the X. We find that under some parameter combinations (Figure 10), RND decreases with increasing effective M-GOUNDRY effective population size, which could result in a smaller RND value on the autosomes since the autosomes should have an effective size at least as big as the X. However, most parameter combinations suggest that this pattern is unexpected (i.e. most regions of the curves predict that RND should increase with increasing effective population size), and the estimate for the ancestral effective size of M-GOUNDRY we obtained in a separate demographic analysis above suggests that these subgroups exist in a parameter space where the RND function is consistently increasing with increasing effective sizes. Taken together with the demographic inference, the above results suggest that, after having initially diverged ecologically approximately 100,000 years ago, genomic barriers to introgression have only been established within the last 100 years, presumably owing to the accumulation and extended effects of locally adapted loci or genetic incompatibility factors within large swept X*h* region on the GOUNDRY X chromosome

**Figure 10:**
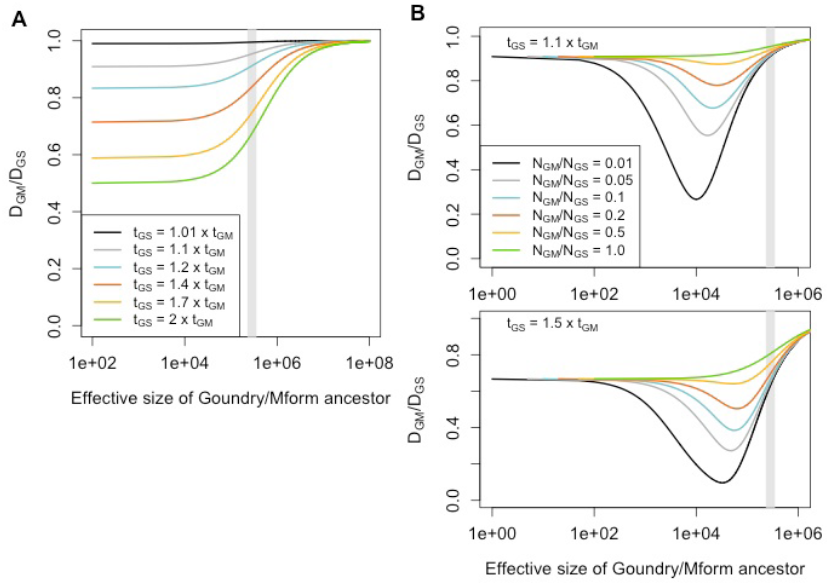
Modeling expected values of Relative Node Depth (*D*_*GM*_/*D*_*GS*_). **A)** Expected values of RND when ancestral population sizes are assumed to be equal. Colors indicate the expectations under different relative split times. **B)** Expected values with t_GS_ split time fixed to 1.1 (top) times the split time between GOUNDRY and M (t_GM_) or 1.5 times (bottom). Colors indicated relative effective sizes of ancestral populations. Values are plotted as a function of the GOUNDRY-M effective size (x-axis). Grey bar indicates 95% confidence interval demographic estimate for GOUNDRY-M ancestral size (see Methods).

Collectively, the data show that nucleotide-diversity corrected divergence is higher inside the putative inverted region, the inverted region as a chromosomal segment is the most diverged of all segments of the same size on the X chromosome, and the X chromosome as a whole is more diverged among GOUNDRY and M form relative to the autosomes. The most parsimonious explanation for these patterns is that, although very few new mutations have accumulated inside of X*h* since its origin less than 100 years ago, ongoing introgression among M form and GOUNDRY has led to a greater density of shared polymorphism and therefore lower sequence divergence in non-inverted regions of the X chromosome relative to the inversion, especially distal to the inversion breakpoints. These results lead us to conclude that while cladogenesis of GOUNDRY and M form ∽100 kya by other means established some degree of temporally fluctuating reproductive isolation, the recently derived putative X*h* inversion now serves as a genomic barrier to introgression, and the effects of selection against migrant haplotypes have begun to extend to linked sites outside the inversion breakpoints.

### Collinear X chromosomes are barriers to introgression

The X chromosomes of the *A. gambiae* M and S forms are collinear with that of *A. merus*, so recombination is not expected to be suppressed in contemporary or historical hybrids among these groups. To test whether sequence divergence is exceptionally high on the X relative to the autosomes in these comparisons despite the lack of inverted regions, we compared genetic divergence and nucleotide diversity among genomic regions as described above. We find that genetic divergence between M and S scales with nucleotide diversity across the genomic regional classes (Figure 9), consistent with few long term barriers to introgression. However, inspection of chromosomal distributions of divergence reveals a slight elevation in genetic divergence in a chromosomal region where nucleotide diversity is low relative to the rest of the chromosome (∽16-20 MB; Figures 4 and 7) and recombination is known to be proportionally rare in M-S hybrids [88]. In contrast, genetic divergence between *A. gambiae* (M form) and *A. merus* does not scale with nucleotide diversity (Figure 9). Nucleotide diversity is lower on the X than on the autosomes (M-W *P* < 2.2×10^−16^), but genetic divergence is significantly higher on the X (M-W *P* < 2.2×10^−16^). Moreover, we find that pericentromeric and telomeric regions are significantly more diverged than the freely recombining autosome (M-W *P* < 2.2×10^−16^), but nucleotide diversity is significantly lower in these regions (M-W *P* < 2.2×10^−16^). Although we could not test for recent introgression among *A. gambiae* and *A. merus* using the ABBA-BABA tests above, this pattern of excess divergence in some genomic regions is in contrast to what we would expect under a divergence in allopatry model and is more consistent with historical introgression in autosomal regions among *A. gambiae* and *A. merus*, likely in the evolutionary period following the initial species split. Overall, these findings are consistent with a large role for the X in driving speciation even among subgroups with collinear X chromosomes and pericentromeric regions diverging secondarily.

## Discussion

We present one of the first analyses of complete genome sequences from multiple species and subgroups from the *Anopheles gambiae* species complex, including the newly discovered cryptic GOUNDRY subgroup of *A. gambiae*, and demonstrate the power of multi-subgroup analysis to comprehensively dissect the mosaic pattern of divergence, adaptation, and introgression across the genome at base-pair resolution. Using average genetic distance across whole genomes, we show clear monophyly of each subgroup and species and confirm that *A. gambiae* is more closely related to *A. arabiensis* than to *A. merus.* We also show that, in contrast to initial findings [17], the newly discovered GOUNDRY subgroup of *A. gambiae* falls genetically within the molecular forms of *A. gambiae* and is not an outgroup. Our demographic analysis suggests that GOUNDRY has existed for approximately 100,000 years and represents a recent example of the frequent speciation dynamics in *Anopheles* that we think is common.

We also show that the genomes of these species and subgroups have been subjected to extensive natural selection in recent evolutionary history, likely related to local adaptation to distinct ecological niches, and we provide the first base-pair resolution maps of recent selective events. We annotated each putative selective event with an associated gene and provide many actionable candidates for follow-up experiments. Interestingly, the selection maps are almost entirely distinct, reflecting ongoing adaptive divergence among these taxa. Despite extensive overlap in their geographic distributions, it is likely that these taxa differ in important ecological and physiological phenotypes [8,14]. For example, recent evidence shows that *A. gambiae* M form and *A. arabiensis* use distinct strategies for surviving the dry season in West Africa and re-establishing local populations after the first rain [89]. In contrast, some ecological pressures are shared among these species as evidenced by the apparent selective benefit of sharing the 2L*a* chromosomal inversion. Moreover, the presence of insecticides like dieldrin has made insecticide resistance alleles globally beneficial among the species studied here. Interestingly, we show that the M form has experienced significantly more positive natural selection than GOUNDRY. Since we know so little about GOUNDRY, it is difficult to speculate on the fundamental differences underlying this discrepancy. One possibility though is that the M form subgroup, having expanded into a presumably much larger and ecologically varied geographic distribution, continues to face new adaptive challenges, while GOUNDRY may exist in a relatively small, specialized and ecologically homogeneous environment with fewer novel selective pressures. Substantial field and functional studies focused on the ecology, mating behavior, and physiology of these subgroups would help to explain the selective pressures underlying their divergence.

We show that the physical locations of putatively introgressed genomic segments coincide with the location of selective sweeps in one or both of the exchanging taxa and suggest that some of these selective events were driven by advantageous alleles shared through introgression. Adaptive introgression has been documented in other systems as well as *Anopheles* [6,62], and appears to be an important evolutionary channel for functional genetic variants that can facilitate phenotypic adaptation. In addition to the compelling case of introgression of a *Rdl* insecticide resistance allele (Figure 8), we identify 17 other candidate sweeps that may have involved an introgressed beneficial allele. We note that identifying selective sweeps and identifying introgressed haplotypes are each challenging tasks in their own right, each with their own confounding issues. Indeed, these two evolutionary processes have the potential to confound the population signal of the other resulting in both false negative and false positive selection scan and introgression results, although the test for selective sweeps used here was shown to be robust to some plausible population genetic scenarios that included both migration and selection [43]. Further exploration is needed to understand fully how these processes interact and develop more sophisticated methods to explicitly test for adaptive introgression.

The nature of species boundaries and divergence among species and subgroups of *Anopheles* is not well understood, despite considerable effort [13,29,30,35,90]. The original observation of centromeric ‘islands of divergence’ between the M and S molecular forms led to the conclusion that gene flow among these species was extensive and ongoing, but these islands remained differentiated for reasons presumably related to their role in speciation among these taxa [29]. Subsequent studies observed especially high levels of differentiation in the ‘islands of divergence’ against a background of differentiation across most of the genome, leading to the conclusion reproductive isolation among these taxa is more advanced than previously thought [13,30]. However, these studies did not determine the relative roles of genetic drift, natural selection on linked sites, and introgression in determining the observed levels of differentiation.

In this study, we use various methods that distinguish between these population genetic processes and provide insight into whether introgression has occurred recently and which genomic regions may be barriers to introgression. We show that chromosomal segments have recently introgressed between the subgroups of *A. gambiae* as well as between *A. gambiae* and *A. arabiensis* (Figure 6). Importantly, our analysis captures the physical locations and genomic coverage of introgressed regions. However, we note that these introgressed haplotypes were identified in a set of representative individuals and therefore likely reflect a combination of stochastic drift processes as well as deterministic selective forces following introgression. The specific composition of introgressed haplotypes along the genome among individuals in each population likely differs to some extent. Over the evolutionary time scales, we expect that natural selection will play a large role relative to drift in determining the composition of ancestry along the genome resulting in a mosaic of introgressed segments along the genome, as has been observed for Neanderthal ancestry in modern humans [25]. To test for heterogeneity in ancestry that may reflect the action of natural selection and therefore barriers to introgression, we measured genetic sequence divergence among taxa and showed that the long-term differential introgression has indeed shaped patterns of divergence among species and subgroups of *A. gambiae* and *A. arabiensis* (Figure 9). Our divergence analysis points to an exceptional history of autosomal introgression among *A. gambiae* and *A. arabiensis* that has created dramatic differences along the genome in genetic affinity among these species (Figure 9). Combined with the evidence that introgression has transferred globally beneficial alleles, such as insecticide resistance alleles, it is clear that introgression has played an important evolutionary role in this species complex and can have large effects on public health.

We show that the X chromosome is the most diverged and harbors the least evidence for introgression, suggesting that it is likely a barrier to introgression in multiple *Anopheles* species (Figures 6 and 9). In some cases, chromosomal inversions seem to play an important role. The observation that the X chromosome plays a disproportionately large effect in driving speciation (‘large-X’) is in line with studies from many organisms ranging from *Drosophila* to mammals [23,25,81,91], but a unifying explanation for this pattern has yet to emerge. Multiple hypotheses have been posited to explain the underlying evolutionary mechanisms underlying this pattern including the ‘faster-X’ hypothesis, sex ratio meiotic drive, and mis-regulation of X-linked genes in males (reviewed in [92]). The ‘faster-X’ hypothesis posits that X-linked loci adapt faster than autosomal loci because X-linked recessive mutations are exposed to selection in males that have only a single X chromosome, resulting in faster accumulation of hybrid sterility factors [23,80]. Another popular hypothesis involves sex ratio meiotic drive where species-specific sex ratio distorter suppressors are disrupted in hybrids causing hybrid sterility [93,94]. A third hypothesis to explain this pattern is that gene expression dosage compensation of X-linked genes is mis-regulated in hybrids, causing sterility in hybrids [95]. Although data have accumulated in *Drosophila* allowing more detailed speculation about the mechanisms underlying the large-X effect [92], more data are needed to fully understand this pattern in *Anopheles*.

Interestingly, we also show that pericentromeric regions also harbor especially high levels of divergence among more distantly related species pairs, suggesting a model where the X chromosome is the first barrier to gene flow and these regions differentiate secondarily. Implicit to this model is that different regions become barriers to introgression in stages while introgression continues to homogenize the remaining genomic regions as in standard speciation with gene flow models [96]. This pattern is most apparent in the comparison between *A. gambiae* and *A. merus* (Figures 7 and 9) and provides strong evidence that both *A. arabiensis* as well as *A. merus* diverged from *A. gambiae* while continuing to hybridize at a non-negligible rate in at least the early stages of speciation. In the case of *A. arabiensis*, we show that contemporary hybridization with *A. gambiae* continues despite strong barriers to genetic introgression across the X chromosome. Similar patterns of elevated divergence in lowly-recombining pericentromeric and telomeric regions have also been observed in comparisons between *Drosophila* species [26,28,97,98], but our results are the first demonstration of this pattern of excess sequence divergence in *Anopheles*. It is important to note that our results, especially the observation of excess divergence between *A. gambiae* and *A. merus* in low-diversity pericentromeric regions, are robust to the issues confounding previous observations of high differentiation in these regions, since our results derive from absolute measures of divergence instead of relative divergence measures that are highly sensitive to other population genetic processes including natural selection [32,33,35].

The ABBA-BABA test has become a preferred method for detecting introgression, but there are several caveats and concerns relating to both the standard test as well as our modified *D*_*BLOCK*_ test. First, it is possible that the signals of introgression we have detected are not from the species used in the test, but in fact we have detected introgression from an un-sampled, or ‘ghost,’ species. Durand et al. [56] showed that such introgression can affect the results of ABBA-BABA tests. The presence of ‘ghost’ *Anopheles* species hybridizing with the species sampled here is certainly a possibility and could impact some of our results. However, the possibility of introgression from ‘ghost taxa’ does not change our conclusion that introgression continues among *Anopheles* species shaping patterns of divergence regardless of exactly which subgroup is the donor. Second, a recent analysis showing that a similar block-wise test of introgression lacked power to identifying introgressed haplotypes [99] raises questions about the robustness of our *D*_*BLOCK*_ analysis. However, there are two important differences between our approach and the one evaluated by Martin et al. [99]. While these authors implemented the test based on constant-sized physical windows of the genome, we used constant-sized blocks of informative sites that varied in their physical size. This is an important distinction because our approach controls for the amount of information in each window, while the number of ancestry informative sites is bound to vary greatly among physical windows in the approach of Martin et al., which is likely to impact the sensitivity of this approach. The second difference between the approaches is that our approach explicitly controlled for linkage disequilibrium in the data. We chose block sizes of 500 informative sites because this resulted in average physical window sizes of ∽ 250 to 350 kb (see Methods), which allowed each block to be divided into 100 segments larger in size than the expected distance that LD decays to background levels in this system. As a result, we believe that our approach is a robust approach for identifying introgressed genomic regions and is not likely to suffer from the same concerns raised by Martin et al. [99].

Our data point to a model where the speciation process is reticulate with offshoot populations and ancestral species remaining connected by periodic hybridization of differential efficiency across the genome that sometimes facilitates adaptation (Figure 11). Although the region of the genome that we identify with statistical evidence for recent introgression is relatively small (∽3%) among the individuals tested here, the effect on the autosome is large over evolutionary time. The relationship between *A. gambiae* and *A. arabiensis* provides an extreme example of variation in the permeability of species boundaries, with evidence of autosomal introgression over the evolutionary history of these two species, including the sharing of an insecticide resistance allele in the last 60 years. In stark contrast to this pattern, the X chromosomes of these species remain highly diverged relative to the autosomes, implying that introgression has been ineffective on the X and that the X chromosome is the primary driver of speciation. A recent independent comparison between the *A. arabiensis* and *A. gambiae* genomes supports our conclusion that introgression is a major factor in the evolutionary history of the *A. gambiae* species complex [100].

**Figure 11:**
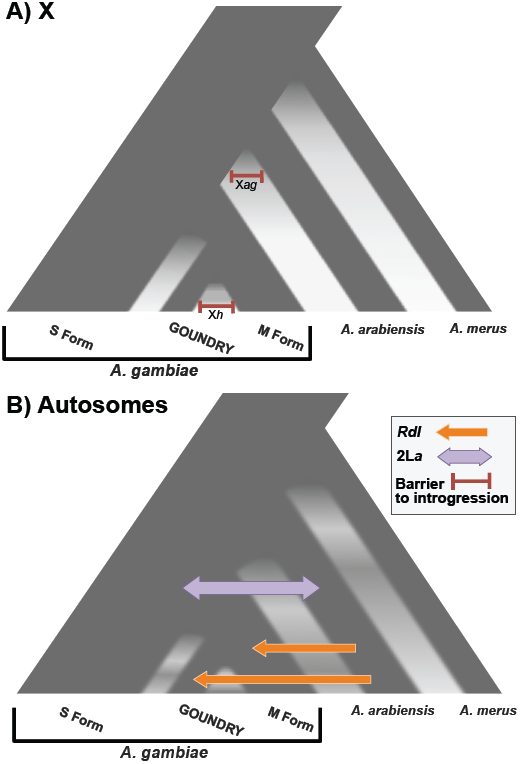
Evolution of founder populations in a species complex. Reproductive isolation among ecologically specialized founder populations and their ancestral population establishes differentially across the genome. **A)** Introgression is ineffective on the X chromosome in general, and chromosomal inversions may serve as barriers in some cases. **B)** Amongst a background of genetic divergence and adaptive differentiation on the autosomes, introgression maintains genetic connectivity across the speciation continuum occasionally allowing rapid adaptation to novel environments in the recipient population, including arid habitats (2L*a*) and insecticides (*Rdl*).

In sum, our results suggest that species and subgroups in the *A. gambiae* species complex comprise a diffuse and interconnected gene pool that confers access to beneficial genetic variants from a broad geographic and environmental range. Such genetic affinity has important implications for malaria control. On one hand, transgenes may spread more easily among subgroups and species of malaria vectors, which could reduce the effort needed to reach and manipulate all populations involved in disease transmission. On the other hand, our analysis suggests that ecological specialization of new subgroups is common and eradicating a population without removing the ecological niche may simply open the niche for a new population to invade with potential a new and unknown consequences for disease transmission. In both cases, our results underscore the complexities involved in vector control on a continental scale.

## Materials and Methods

### Mosquito samples

Mosquito sample collection and species/subgroup identification was previously described for the M form (recently re-named *A. coluzzii* [8]), GOUNDRY, and *A. arabiensis* samples [17]. Briefly, larvae and adults were collected from three villages in Burkina Faso in 2007 and 2008 (Table S5). Larvae were reared to adults in an insectary, and both field caught adults and reared adults were harvested and stored for DNA collection. One *A. gambiae* S form individual was also included in this study. This sample was collected indoors as an adult in the village of Korabo in the Kissidougou prefecture in Guinea in October 2012. Individuals were typed for species, molecular form and *2*L*a* karyotype using a series of standard molecular diagnostics [101–103]. All M form and *A. arabiensis* samples are *2*L*a^a/a^* homokaryotypes and the S form sample typed as a heterokaryotype (*2*L*a^a/+^*). Eleven of the twelve GOUNDRY samples typed as *2*L*a^+/+^* homokaryotype, but one sample (GOUND_0446) typed as a *2*L*a^a/a^* homokaryotype.

### DNA extractions and genome sequencing

DNA was extracted from female carcasses using standard protocols. Genomic DNA samples were diluted in purified water and sent to BGI (Shenzhen, China) for paired-end sequencing on the Illumina HiSeq2000 platform. We sequenced 12 females from the GOUNDRY subgroup of *A. gambiae*, 10 females for the M form subgroup of *A. gambiae*, and 9 female *Anopheles arabiensis*. Sequence reads were filtered if they 1) contained more than 5% Ns or polyA structure, 2) contained 20% or more low quality (Q<20) bases, 3) contained adapter sequence, or 4) contained overlap between pairs. After quality filtering, each sample was represented by an average of 47.91 million reads with an expected insert size of 500 base pairs, except two samples that were sequence to deeper read depth (Table S1).

In a separate sequencing effort, we also sequenced one individual *A. gambiae* S form female. The sequencing library was prepared using the Nextera kit (Illumina Inc., San Diego, CA) according to manufacturer’s specifications and sequenced on the Illumina HiSeq2000 platform (Illumina Inc., San Diego, CA) at the University of Minnesota Genomics Center core facility for a total of 125.88 million reads.

All Illumina reads newly generated for this study have been submitted to the Short Read Archive at NCBI under accession IDs ranging from XXXXXX-XXXXX.

We obtained publicly available *Anopheles merus* and *Anopheles arabiensis* short read Illumina data from NCBI SRA. We downloaded *A. merus* accessions ERR022713-8 that were generated and deposited by The Sanger Center. Individual accessions are paired-end 76 bp read fastq files representing whole genome sequence from single individual *A. merus* individuals collected Kenya sequenced on the Illumina GAII platform. In addition, we downloaded *A. arabiensis* accession SRX377561, representing an individual collected in Minepa, Tanzania [38]. These data are paired-end 100 bp reads generated on the Illumina HiSeq2000 platform. This sample is the individual sequenced to ‘high coverage’ from the Marsden et al. Tanzania population sample.

### Illumina short read alignment and subgroup specific references

At the time of this analysis, the only genome reference sequence with scaffolds mapped to chromosomes for any *Anopheles* mosquito was the *Anopheles gambiae* PEST AgamP3 assembly [[104]; vectorbase.org], so we used this reference for mapping reads from all groups and species in this study. Since an unknown subset of the short reads generated for this experiment were expected to be substantially diverged from the PEST reference, we conducted short read alignment iteratively in two steps. First, we conducted paired-end mapping of all available reads to the *Anopheles gambiae* PEST reference genome [104] using the BWA *mem* algorithm [[105]; bio-bwa.sourceforge.net] with default settings except applying the –M flag, which marks shorter hits as secondary. We then generated new ‘Subgroup-Specific’ references (hereafter SPEC) for *Anopheles gambie* M form, *Anopheles gambiae* GOUNDRY, *Anopheles arabiensis*, and *Anopheles merus* separately. To do so, we combined sequence reads for each Subgroup and generated a pseudo-consensus sequence for each Subgroup. We used the software package ANGSD (version 0.534; [44]; popgen.dk/wiki) to generate a read pileup, count reads and bases at each site, and identify the major allele at all sites. Then for every site that was covered by 1) at least 4 sequence bases that pass filtering, 2) fewer than a Subgroup-specific threshold number reads (mean read depth plus two standard deviations), and 3) a major allele that was segregating at a frequency of at least 0.5 across all reads, we assigned the new reference base to the major allele and to ‘N’ otherwise.

After alignment to the PEST reference, the mean total read depth for the autosomal arms was 136.14 for *A. arabiensis*, 136.63 for *A. gambiae* M form, 157.42 for *A. gambiae* GOUNDRY, and 52.94 for *A. merus*. Mean total read depth on the X chromosome was 124.27 for *A. arabiensis*, 145.63 for *A. gambiae* M form, and 168.75 for *A. gambiae* GOUNDRY, and 44.37 for *A. merus*. The proportion of unknown bases (‘N’s) increased in the new references (0.12, 0.11, 0.05, 0.14 for the autosomes of *A. arabiensis*, M form, GOUNDRY, and *A. merus*, respectively, and 0.22, 0.05, 0.06, 0.25 for the X) relative to PEST (0.02 for autosomes, 0.04 for X). The differences between Subgroups in read depth and proportion of sites missing data can be attributed in part to differences in the number of individuals sequenced (n = 9,10,12, 6 for *A. arabiensis*, M form, GOUNDRY, *A. merus*, respectively). However, the differences between autosomes and the X likely reflect difficulties in mapping divergent reads to the PEST reference. The proportion of sites missing data were higher on the X relative to the autosomes for alignments of data from *A. arabiensis* and *A. merus* to both the PEST reference and the Subgroup specific references (Table S6). Two other studies found a similar discrepancy when mapping these species to the *A. gambiae* PEST reference [37,38], indicating that it is not unique to our pipeline and likely reflects the relatively higher divergence on the X that proves to be a higher barrier to read mapping.

Following the generation of new reference sequences for *A. gambiae* M form, GOUNDRY, *A. arabiensis,* and *A. merus*, short read datasets for each Subgroup were then aligned back to the new subgroup specific (hereafter SPEC) reference using the BWA *mem* algorithm with default settings. The S form individual was aligned to the M form reference sequence. Local realignment around indels was then performed for each subgroup separately using GATK [106]. Duplicates were removed using the SAMtools *rmdup* function.

Although the read mapping rate was consistently lower for the SPEC reference relative to rates when mapping to PEST, the proportion of mapped reads with a quality score of 20 increased for all groups except the M form Subgroup where it decreased slightly. The reduction in mapping rate when using the SPEC reference reflects the greater number of ambiguous bases (‘N’) in the SPEC reference relative to PEST and thus a smaller mapping target. Importantly, the proportion of mapped reads with quality score of 20 on the X chromosome increased from 0.6961 (PEST) to 0.7775 (SPEC) when mapping the *A. merus* data to the PEST and SPEC references. A similar increase was also observed for *A. arabiensis*, with the proportion of Q20 reads increasing from 0.7604 (PEST) to 0.8241 (SPEC), indicating that the Subgroup specific reference is especially helpful for mapping X-chromosome reads. As discussed above, mapping non-*A. gambiae* reads to the PEST X chromosome is expected to be difficult. Mapping biases may lead to underestimates of diversity and genetic divergence, but such downward biases do not change the main conclusions regarding *A. arabiensis* and *A. merus*.

### Data Filtering

The data were filtered in two steps. First, we used SAMtools to generate a pileup and genotype likelihoods for each site. We calculated a series of alignment and read statistics and then applied a series of filters to obtain a set of sites considered reliable for downstream analysis. Filters were applied using the SNPcleaner Perl Script from the ngsTools package [46]. Those filters are as follows.

1. *Read distribution among individuals*: No more than one individual is allowed to have fewer than 2 reads covering the site.
2. *Maximum read depth*: The site must not be covered by more than 350, 350, and 400 reads for M form, *A. arabiensis*, and GOUNDRY, respectively, in each combined sample.
3. *Mapping Quality*: Only reads with a BWA mapping quality of at least 10 were included.
4. *Base Quality*: Only bases with Illumina base quality of 20 or more were included.
5. *Proper pairs*: Only reads the mapped in the proper paired-end orientation and within the expected distribution of insert lengths were included.
6. *Hardy*-*Weinberg proportions*: Expected genotype frequencies were calculated for each variable site based on allele frequencies based on genotypes called with SAMtools. Any site with an excess of heterozygotes were considered potential mapping errors and excluded.
7. *Heterozygous biases*: Sites with heterozygous genotype calls were evaluated with SAMtools for several biases using exact tests. If one of the two alternative alleles was biased with respect to the read base quality (minimum *P*=1×10^−100^), read strand (minimum *P*=1×10^−4^), or distance from the end of the read (minimum *P*=1×10^−4^), the site was excluded.

The next filter was intended to exclude regions of the genome where short-read alignment may be compromised, such as low-complexity and regions with a high concentration of ‘N’ bases in the reference. The goal was to avoid edge effects and regions where mapping becomes ambiguous. To identify such regions, we scanned each Subgroup specific reference calculating the proportion of Ns in every 100-basepair window. All sites that fell within windows that contained 50 or more Ns were excluded. Since short read alignment is likely to be unreliable in highly repetitive genomic regions such as heterochromatic regions, we also excluded regions that have been identified as heterochromatic in *A. gambiae*, including both pericentric and intercalary heterochromatic regions [108] to be conservative in our analyses.

A final list of sites-to-exclude was compiled for each subgroup-specific reference that included any site excluded by any of the above filters. This resulted in a different number and set of sites available for downstream analysis for each Subgroup (Table S7). In addition to these excluded sites, we identified a series of regions that exhibited exceptionally high nucleotide diversity even after front-end filtering. These sites were not excluded from ANGSD analyses (see below), but were excluded from all population genetic analyses (Table S7).

### Site-frequency spectrum inference

Population genetic inference from next-generation sequencing data was performed using a statistical framework implemented in the software package Analysis of Next-Generation Sequencing Data (ANGSD, [44], popgen.dk/wiki). Much of the BAM manipulation and read-filtering functionality implemented in SAMtools is also implemented in ANGSD and several functions were utilized in all of the following analyses with ANGSD. Minimum map quality and base quality thresholds of 10 and 20 were used. Probabilistic realignment for the computation of base alignment quality (BAQ) was enabled throughout with a downgrading coefficient of 50. Reads that were not aligned as proper pairs or had bitwise flags above 255 were excluded.

The first step in the pipeline was to infer the site-frequency spectrum (SFS) directly from genotype likelihoods estimated from the read data. The global SFS was estimated in two steps. The first step was to obtain a maximum likelihood estimate of per-site allele frequencies by calculating multi-sample genotype likelihoods using the SAMtools model ([107]; -GL 1 in ANGSD) and estimating the minor allele frequency using the –realSFS 1 function in ANGSD [44]. Files containing the per-site estimates of allele frequencies were then used as input for optimization of the global SFS across all sites using a BFGS optimization algorithm implemented in the optimSFS program within ANGSD. Unfolded spectra were obtained by including ancestral polarization assigned by a synthetic ancestral sequence described below. Sites without ancestral assignment were excluded from SFS estimation. The optimized global SFS was estimated for each chromosomal arm separately and the results from 2R, 3L, and 3R were averaged to obtain an autosomal SFS. Autosomal and X chromosome site frequency spectra were obtained for M form, GOUNDRY, and *A. arabiensis* separately. These spectra were used as priors for downstream analyses within ANGSD and are shown in Figure S6.

### Ancestral sequence

We generated a synthetic ancestral sequence by assigning ancestral and derived alleles based on their presence in our M form, GOUNDRY, *A. arabiensis*, and *A. merus* population samples. We first identified all alleles segregating at every site in the genome of each group by calculating genotype likelihoods (-GL 1) and estimating the minor allele frequency (–doMaf 2 in ANGSD). We used all variable sites except singletons. If all groups shared the same allele at a site or if all groups were missing data at a site, the conserved allele or an N was included in the new ancestral sequence, respectively. If some groups lacked data at a site and only one allele was observed in the remaining groups with data, this allele was included in the new sequence. Lastly, if an allele was found to be segregating in at least three of the four groups and the alternate allele was found in only one or two groups, the major allele was chosen as the ancestral allele. This approach is conservative but also flexible in that it requires an allele to be found in either 1) three different species or 2) both subgroups of *A. gambiae* and one outgroup species. It is possible that gene flow among these groups could introduce bias here, but it is difficult to distinguish gene flow from lineage sorting at this stage in the analysis. We compared the SFS spectra inferred using this ancestral sequence to those obtained using *merus* as the ancestral sequence and found a larger enrichment of high frequency derived alleles in the *A. merus-*derived spectra as would be expected if ancestral mis-specification were common (not shown).

### Genotype calling

Genotypes were called in two steps at all sites that passed filtering. For both the population samples (M form, GOUNDRY, *A. arabiensis, A. merus*) as well as the *A. gambiae* S form individual, genotype likelihoods were calculated using the SAMtools model as described above. Then genotype posterior probabilities were calculated for each individual at each site. For the population samples, posterior probabilities were calculated using maximum likelihood estimates of allele frequencies as a prior [109]. For the single individual samples, a uniform prior was assumed for calculation of the posterior probabilities. For all samples, diploid genotypes were called only at sites with posterior probabilities of 0.9 or greater. Genotypes for GOUNDRY sample were obtained using a slightly different approach since this Subgroup is partially inbred (see below).

### Inbreeding analysis

#### Estimating inbreeding coefficients

Initial estimates of the global site frequency spectrum (SFS) in GOUNDRY produced distributions of allele frequencies that deviated substantially from standard equilibrium expectations as well as from those observed in the *A. gambiae* M form and *A. arabiensis* groups. Most notably, the proportion of doubletons was nearly equal to that of singletons in *A. gambiae* GOUNDRY. This observation is consistent with widespread inbreeding in the GOUNDRY subgroup. We tested this hypothesis in two ways, with the goals of both characterizing the pattern of inbreeding in this subgroup as well as obtaining inbreeding coefficients for each individual that could then be used as priors for an inbreeding-aware genotype-calling algorithm. We used the method of Vieira et al. [110], which estimates inbreeding coefficients in a probabilistic framework taking uncertainty of genotype calling into account. This approach is implemented in a program called ngsF (github.com/fgvieira/ngsF). ngsF estimates inbreeding coefficients for all individuals in the sample jointly with the allele frequencies in each site using an Expectation-Maximization (EM) algorithm [110]. We estimated minor allele frequencies at each site (-doMaf 1) and defined sites as variable if their minor allele frequency was estimated to be significantly different from zero using a minimum log likelihood ratio statistic of 24, which corresponds approximately to a *P* value of 10^−6^. Genotype likelihoods were calculated at variable sites and used as input into ngsF using default settings. For comparison, we estimated inbreeding coefficients for *A. gambiae* M form, GOUNDRY, and *A. arabiensis* using data from each chromosomal arm separately (Figure S7).

We inspected the distribution of homozygous regions likely related to inbreeding dynamics and found that these regions are randomly distributed and different among individuals. A more thorough analysis and discussion of these patterns, possible causes, and possible implications will be presented in a separate manuscript.

#### Recalibrating the site-frequency spectrum and genotype calls

We used the inbreeding coefficients obtained above for the GOUNDRY sample as priors to obtain a second set of inbreeding-aware genotype calls and an updated global SFS. We used ANGSD to make genotype calls as described above. However, in this case, we used the –indF flag within ANGSD, which takes individual inbreeding coefficients as priors instead of the global SFS [110]. Similarly, we used the inferred inbreeding coefficients to obtain an inbreeding-aware global SFS. We estimated the global SFS from genotype probabilities using –realSFS 2 in ANGSD, which is identical to –realSFS 1 [44] except that it uses inbreeding coefficients as priors for calculations of posterior probabilities [110].

### Genetic divergence and neighbor-joining tree

#### D_xy_ calculations

We measured absolute genetic divergence as the average number of pairwise differences between alleles from different subgroups, or *D*_*xy*_ [111]. We calculated *D*_*xy*_ as

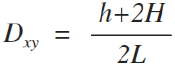

where *h* is the number of sites where one or both individuals carry heterozygous genotypes, *H* is the number of sites where the two individuals are homozygous for different alleles, and *L* is the number of sites where both individuals have called genotypes. As with our estimates of within-population diversity, we excluded all sites within 200 bp of the outer coordinates of annotated protein coding sequences in the AgamP3.8 gene set (vectorbase.org). For subgroups where we have sequences from multiple individuals (M form, GOUNDRY, *A. arabiensis, A. merus*), we estimated *D*_*xy*_ calculated from the number of pairwise differences between each individual from every other subgroup and then averaged across comparisons for each subgroup to obtain an average *D*_*xy*_ value for each pair of subgroups. For comparisons made to the S form, *D*_*xy*_ was calculated between the individual S form sequence and the individual sequence with the highest average read depth from each of the other subgroups.

For chromosomal plots (Figure 7) and boxplot analysis (Figure 9), each chromosomal arm was divided into 10 kb windows and *D*_*xy*_ was calculated across each window. Windows with fewer than 200 sites with data for comparison were excluded. Chromosomal curves were LOESS-smoothed using a span of 1% and degree of 2 with the loess.smooth function in R [112]. For the boxplot analysis, chromosome 2L was excluded to avoid the large effects of the *2*L*a* inversion. Comparisons were made between categories using a Mann-Whitney test implemented in the wilcox.test function in R [112].

#### Neighbor-joining trees

We visualized average genetic relatedness across the genome among individuals and species using neighbor-joining distance trees based on pairwise values of *D*_*xy*_. We calculated *D*_*xy*_ across chromosomes 2R, 3R, 3L, and X, but excluded 2L to avoid the effects of the 2L*a* inversion. *D*_*xy*_ was calculated across intergenic sites at least 200 bp from protein coding sequences in euchromatic genomic regions. Neighbor-joining trees were generated using the *ape* [113] package in R and drawn using the Geneious software (www.geneious.com). We used a bootstrap analysis to estimate confidence in the major, internal node topology (i.e. species relationships) by dividing the dataset into 1 MB physical windows and sampling with replacement 1,000 new datasets. Distance neighbor-joining trees were generated for each bootstrap replicate and compared to the tree made from the true data to determine consistency of major clade placement.

### Demographic inference

#### 2D-spectra and model fitting

We fit population historical models to the two-dimensional site frequency spectrum for GOUNDRY and M form subgroups using the program *dadi* [42]. We obtained the unfolded 2D spectrum for chromosomal arms 2R, 3R and 3L by first estimating allele frequencies at each site for M form and GOUNDRY separately, then calculating posterior probabilities of allele frequencies at each site using the global SFS priors, and finally calculating the 2D spectrum. Posterior probabilities were calculated using the program sfstools distributed with ANGSD. The program 2Dsfs included in the software package ngsTools (https://github.com/mfumagalli/ngsTools; [46]) was used to calculate the 2D spectra with -maxlike set to 1. Per-site allele frequencies were estimated for each subgroup as described above, except GOUNDRY was handled differently to accommodate the inbreeding signal. Since GOUNDRY is partially inbred, we randomly sampled one allele at each position in the genome to be included for analysis thereby eliminating the effects of inbreeding within a diploid genome on estimates of allele frequency. This resulted in a sample size of 12 chromosomes for GOUNDRY. Initial testing suggested that optimization in *dadi* performed better when sample sizes were equal, so we projected the M form sample from 20 chromosomes down to 12 to make the spectra symmetric using the project function with *dadi*. As for before, all heterochromatic regions and low-complexity regions were excluded. We also excluded all sites within 200 bp of exons annotates in the *A. gambiae* PEST AgamP3 gene set. We calculated the 2D spectrum for each of the chromosomal arms separately and summed them to represent the autosome (Figure 2). Chromosome 2L was excludee to avoid the effects of the large segregating 2L*a* inversion. The final spectrum contained 61,254,546 sites, 3,189,751 of which were variable in our samples.

We defined the demographic function to include three epochs (Figure 2). As with all models in *dadi*, the model begins with an ancestral population and proceeds forward in time. In the first epoch, the ancestral population is split into two populations. Each of the three epochs is defined by five parameters: 1) the duration of the epoch, 2) the effective size of M form, 3) the effective size of GOUNDRY, 4) the rate of migration from GOUNDRY into M form, and 5) the rate of migration from M form into GOUNDRY. We defined the demographic function using the following syntax:

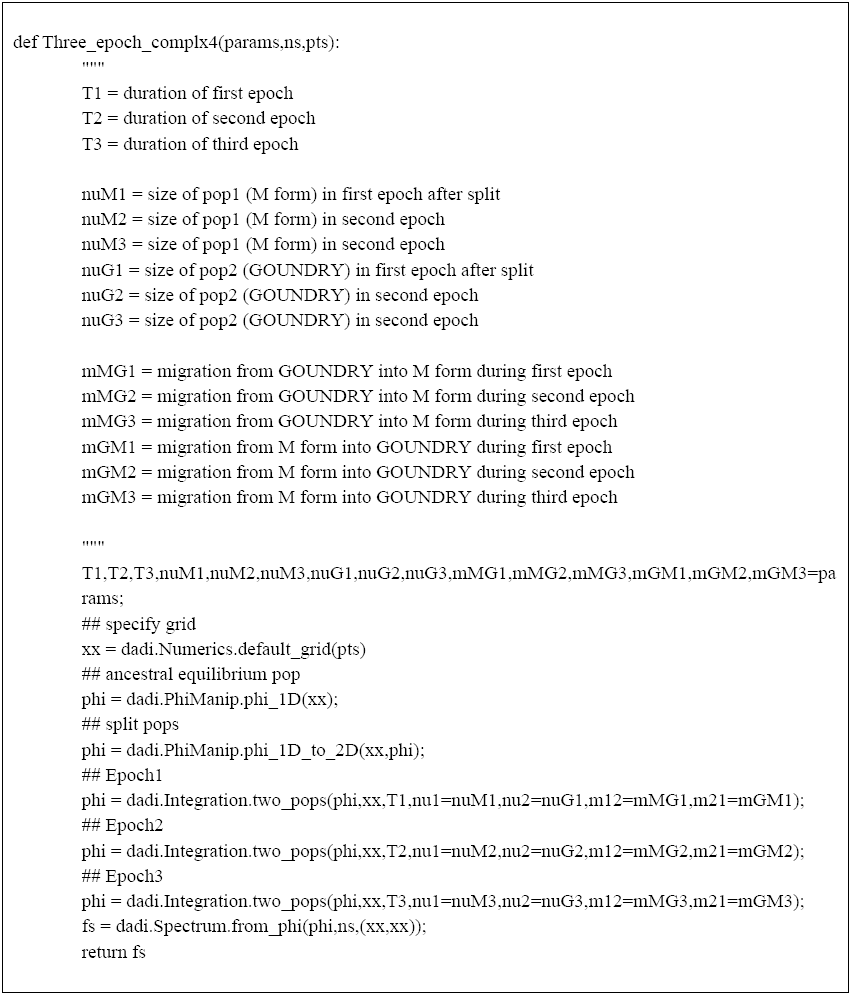

We used the multinomial likelihood approach for evaluating the fit between the data and the 2D spectrum predicted by the model, and conducted inference in two steps. First, we set large boundaries for each parameter and conducted 466 replicate optimizations, perturbing the starting parameters for each run by a factor of 1.5 and allowing 300 iterations per optimization. This step allowed us to manually evaluate the likelihood surface for each parameter and identify regions of the parameter space with high likelihoods for further optimization. Second, to ensure that the optimizations reached best-fit parameter values, we narrowed the parameter search boundaries and conducted a second set of 1,016 optimizations, perturbing the starting values by a factor of 0.5 and allowing 750 iterations. Models that include strong bottlenecks are extremely slow to optimize and not likely to be biologically plausible for systems like mosquitoes with typically large census sizes, so we set lower boundaries of 0.01 for all subgroups sizes for computational efficiency. After ensuring that the optimized values were not hitting boundaries and parameters were converging over replicate optimizations, we chose the optimization with the highest likelihood as the point estimate. To obtain an estimate of the ancestral effective population size, we estimated the population scaled mutation rate (4*N*_*A*_*μ*) that fits the data best using the optimal_sfs_scaling function in *dadi* and solved for *N*_*A*_, assuming a mutation rate of *μ***=** 7.0 × 10^−9^. The spontaneous mutation rate for mosquitoes has not been estimated to date, so we used a mutation rate estimated from *Drosophila melanogaster* as our closest approximation for the mutation rate in *Anopheles*. Multiple estimates of the spontaneous mutation rate for *D. melanogaster* are available from studies differing in their approach, and the estimates vary by an order of magnitude from 2.8×10^−9^ to 1.1×10^−8^ [114–118]. We chose an intermediate value of 7.0×10^−9^.

#### Parameter confidence intervals

We estimated confidence intervals for each parameter using a nonparametric bootstrap approach. We generated bootstrapped genomes by first concatenating chromosomal arms 2R, 3R, and 3L, and then dividing this ‘genome’ into physical 200 kb regions. These regions were sampled with replacement to obtain 100 new genomes with lengths equal to that of the true genome. Since repeating the two-step optimization described above would be computationally infeasible, we conducted optimizations for each bootstrap replicate with a different approach. For each replicate, we conducted 100 optimizations with wide boundaries, allowing 750 iterations and perturbing starting values by a factor of 0.5. We used the maximum likelihood values obtained for the true genome as pre-perturbation starting values. Then for each bootstrap replicate, we chose the optimization with the maximum likelihood. Approximate 95% confidence intervals were calculated for each model parameter as the mean of all replicate values +/-1.96 standard deviations of all replicate values. All bootstrap optimizations were run on the Stampede compute cluster at University of Texas, Austin through the support of the XSEDE program. Optimized parameter values and confidence intervals are reported in Table 1.

### Fixed differences

We identified fixed differences between M form and GOUNDRY using the following criteria. A substitution was inferred if

1. At least (n-2) individuals were represented at a site.
2. The site was considered monomorphic based on allele frequencies inferred using ANGSD being either less than 1/2n or greater than (1-(1/2n)).
3. GOUNDRY and M form harbor different major alleles.

### Linkage disequilibrium

We measured LD using Haploview [119] applied to genotype and SNP calls made using ANGSD as described above for M form, GOUNDRY, and *A. arabiensis*. LD was calculated independently on each chromosomal arm and comparisons were made only between SNPs separated by 10 kb or less. For all analyses, we used the *r*^2^ statistic generated by Haploview. For computational tractability, we reduced the number of SNP-by-SNP comparisons by randomly sampling comparisons down to either 1%, 10% or 20% of the total number of comparisons depending on the total number of comparisons for each subgroup. To obtain LD decay curves, we binned *r*^2^ values for each subgroup based on physical distance in the PEST reference, and then averaged within each bin and plotted as a function of physical distance (Figure S8). We combined measures for chromosomal arms 2R, 3R, and 3L to obtain an autosomal curve while avoiding the effects of the 2L*a* inversion. We estimated a curve for the X chromosome separately. In all three subgroups, LD decays to background levels within several hundred basepairs on both the autosomes and on the X, consistent with previous analyses in these species [38,120,121]. Interestingly, background levels of LD differ substantially among subgroups. Background LD is slightly higher in *A. arabiensis* than in M form, which is consistent with a small effective population size and perhaps some greater degree of population substructure in *A. arabiensis* than in M form. GOUNDRY exhibits very high levels of background LD, but this is due to the high levels of inbreeding-related homozygosity. In all cases, LD decay curves suggest bootstrapping can be conducted on 200 kb regions with confidence.

We generated background LD chromosomal plots (Figure S2), by dividing each chromosome into 10kb windows, identifying all SNPs inside each window, and taking the average *r^2^* value for all comparisons after thinning (see above) between SNPs inside the window and SNPs greater than 1 kb but less than 10 kb away on the PEST reference sequence. Only windows with at least 100 comparisons were included. Background LD was measured for M form and *A. arabiensis* only since inbreeding has skewed measures of LD in GOUNDRY. LOESS-smoothed curves were generated using a span of 1% and a degree of 2 in the loess.smooth function in R [112].

### Scan for recent positive selection

#### Genome scans

We conducted full genome scans for recent complete selective sweeps in *A. gambiae* M form and GOUNDRY as well as *A. arabiensis* using SweepFinder, an implementation of the composite-likelihood test that compares the likelihood of allele frequencies and their physical distribution along a chromosome under a sweep model to the likelihood of the data, given a neutral spectrum of allele frequencies [43]. To assemble input data, we estimated unfolded allele frequencies from genotype likelihoods using the method of Kim ([109]; -doMafs 5 in ANGSD) with ancestral state assigned based on the ancestral sequence constructed as described above. We converted ANGSD output to SweepFinder input files and ran SweepFinder using a grid size that corresponds starting points every 1kb. Instead of using SweepFinder to estimate the global site frequency spectrum, we provided SweepFinder with spectra estimated independently using realSFS in describing SFS inference above.

#### Neutral simulations in GOUNDRY and M form

To establish critical thresholds for test statistics, we simulated population samples of 50 kb M form and GOUNDRY haplotypes using coalescent simulations under the demographic model estimated using *dadi* (described above). We used the scaled mutation rate, *θ*, estimated in the *dadi* inference. Recombination rates are not known in this system, so we conducted a limited set of coalescent simulations under a range of scaled recombination rates. LD decay curves from simulated datasets were compared to LD decay curves in the M form data. We found that increasing the scaled recombination rate by a factor of 5, 6, or 7 relative to the scaled mutation rate produced LD decay curves comparable to the M form data. These factors correspond to per site scaled recombination rates of 0.0177, 0.0212, and 0.0247 compared to a scaled mutation rate of 0.0035. To allow for some variation in recombination rate, we used a mixed distribution of scaled recombination rates with probability of 0.5, 0.25, and 0.25 for 0.0177, 0.0212, and 0.0247, respectively. We conducted all simulations using MaCS ([122]; version 0.5d) with –h equal to 1 to take full advantage of the Markovian approximation of the coalescent implemented in this method. Consistent with the analytical correspondence between the approximate coalescent in MaCS and the full coalescent, simulations have shown that data generated under this approximation compare quite well to data generated under standard full coalescent models [122]. Accordingly, the two dimensional site frequency spectrum calculated from the simulated data closely resembles both the spectra from our data and the maximum likelihood model inferred by *dadi* (Figure S9). The LD decay curves from the simulated data also follow a similar decay rate as the real data (Figure S9).

We used the following MaCS command for autosomal loci:

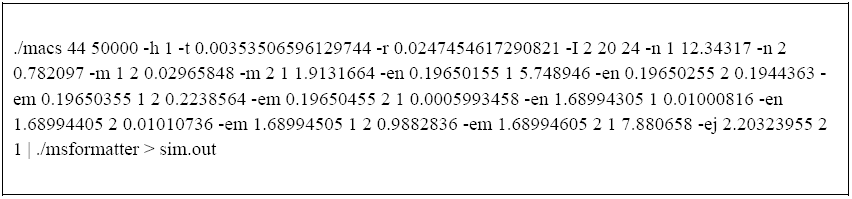

For autosomal loci, we simulated 792,800 50 kb regions corresponding to 200 complete autosomes and applied SweepFinder to the M form and GOUNDRY simulated haplotypes separately. We used a grid size of 50 for SweepFinder to be consistent with the 1kb scale used for the real data. The background site frequency spectrum was estimated by SweepFinder for each 50kb region and used for that region. To simulate data resembling the X chromosome, we simulated 150,000 50 Kb regions under the same demographic model, but applied a ¾ correction to the ancestral effective size inferred from *dadi* and adjusted simulation parameters accordingly.

To determine critical thresholds for a 50 Kb region (hereafter referred to as the per-locus threshold), we found the maximum log likelihood ratio (LLR) found in each 50 kb region for M form and GOUNDRY separately (Figure S10). Then, to obtain per-locus *P* values, the maximum LLRs for all M form and GOUNDRY selective sweep windows in the real data were compared to the corresponding distribution of LLRs calculated from simulated regions. To obtain a genome wide critical threshold corrected for multiple testing, we assembled autosomal 50 Kb regions and X-linked 50 kb regions into synthetic 218.23 Mb genomes and identified the highest LLR within each simulated genome (Figure S10). After identifying windows with LLR values significant at the genome level, we then identified the highest point in each string of continuous significant windows to find the ‘peak of the peak’.

We found that peaks were unexpectedly clustered across the genome, suggesting that some peaks may be fractured evidence of the same selective event. Since SweepFinder has substantially more power when monomorphic sites are included in the analysis [123], we re-ran SweepFinder using both monomorphic and variable sites within clustered regions. We identified clusters of peaks where significant peaks were found within 100 Kb of each. To determine whether multiple peaks separated by windows with non-significant LLR values were actually part of a single large peak, we asked whether more than four continuous windows with LLR values less than 20 separated the peaks in the ‘all-sites’ analysis. SweepFinder is prohibitively slow when monomorphic sites are included in the analysis, so we could not conduct this version across the whole genome. In some cases, this high-resolution analysis revealed a single continuous peak. We present one representative example in Figure S11 where non-significant windows in the variable-site-only analysis separate a number of significant peaks, but the all-site analysis includes only a single low-LLR window separating the peaks. To be conservative, clusters of peaks satisfying these criteria were collapsed into a single peak and the window with the highest LLR was used for annotation. In other cases, however, the all-sites analysis did not provide evidence a single continuous peak, despite an overall peaked shape of the significant windows in the region (Figure S11). The final set of selective sweeps after collapsing clusters is presented in Table S2.

#### More selection in GOUNDRY or M form?

Selective events may not always be identified as single events by SweepFinder and thus our count of independent selective events may be biased, we chose to quantify the proportion of megabases (n = 230) in each genome that harbors at least one selective sweep (i.e. one significant window). To determine whether the difference between M and GOUNDRY is statistically significant, we randomly permuted subgroup assignments of each selective sweep in our data 10^4^ times and conducted the same analysis. We could not make the same comparison with *A. arabiensis* because a different method was used to identify significant signals of selection for that species.

#### Identifying peaks in *A. arabiensis*

We did not estimate a demographic model for *A. arabiensis*, so we used a different approach to identify credible signals of recent selective sweeps. We searched the LLR surface for peaks where the peak included at least two adjacent windows with LLRs in the top 0.1% genome wide. The 0.1% threshold corresponds to a value of 15.88 in this dataset. After finding all such cases, we used the window with the highest LLR in the cluster to represent the peak. This approach identifies 34 distinct peaks across the autosomes and none on the X chromosome.

#### Annotating peaks

For each selective sweep, we found the protein coding sequence most closely associated with the highest point on the LLR peak. If the maximum LLR window fell within a gene in the Agam3.8 gene set, information from this gene was included in the annotation database. If the maximum LLR window fell outside known genes, information from the gene with the nearest 3’ or 5’ boundary was used for the peak. To annotate each sweep, we downloaded information from Vectorbase.org for AgamP3.8 gene set from the *A. gambiae* PEST genome, including gene names, membership in ImmunoDB [124], best GO annotations, and best KOG annotations. Information for all selective sweeps can be found in Tables S2 and S3.

### Nucleotide diversity

To enrich our data set for neutrally evolving sequence, we excluded sites within 200 bp of a coding sequence annotated in the AgamP3.8 gene set for the *A. gambiae* PEST reference available on VectorBase.org from our estimates of nucleotide diversity. If only a single individual was available for a group (*A. gambiae* S form), nucleotide diversity was estimated simply as the fraction of sites with genotypes that were called as heterozygous. If a population sample was available (GOUNDRY, M form and *A. arabiensis, A. merus*), we estimated nucleotide diversity directly from the read data using a maximum likelihood approach based on posterior probabilities of per-site allele frequencies [45]. The method is implemented under the –doThetas function within ANGSD and takes a global SFS as a prior to calculate posterior probabilities of allele frequencies. We chose to use Tajima’s π statistic [125] as our estimate for nucleotide diversity. For all downstream analyses, nucleotide diversity was calculated as an average within 10 kb non-overlapping windows. The comparison of nucleotide diversity between genomic regions (see below) included only 10 kb windows containing at least 500 sites from 2R, 3L, 3R, and X chromosomal arms. The distribution of nucleotide diversity was compared between genomic regions using a Mann-Whitney test implemented in the wilcox.test function in R [112].

### Introgression analysis and D statistics

We used permutation and jackknife analyses to conduct two ABBA-BABA-based tests for introgression. For the set of tests, we divided the genome into blocks of 500 informative sites (i.e. ABBAs and BABAs). These genomic blocks were then divided into 100 segments of 5 informative sites. We chose this block size because this number of sites corresponded to a physical size of ∽100 × L_LD_, where L_LD_ is the physical distance at which point linkage disequilibrium decays to background levels. As a result, we could then divide the genomic blocks into 100 segments that would be larger in physical size than the L_LD_. Since introgressed haplotypes will be highly correlated over exceptionally high physical distances relative to ancestral haplotypes, we expect that the size of introgressed haplotypes would exceed L_LD_ while ancestral haplotypes would not. For the tests involving S form and *A. arabiensis* as the H3 taxon, the mean genomic block length was ∽250 Kb and ∽350 Kb, respectively. Since L_LD_ is approximately 200 bp in this system (Figure S8), and segments within the blocks were approximately 2.5 or 3.5 Kb in size, segments exceeded L_LD_ by a factor of more than 10x. These analyses were conducted using pseudo-haploidized genomes from the individuals from each population/species with the highest short-read coverage. In general, sites involved in calculating *D* are not entirely independent due to linkage disequilibrium, so permutation and jackknife analyses are necessary to properly test for significance. We have used these corrected tests for significance in the following ways.

For the first test, we calculated the *D* statistic calculated for each genomic block (hereafter *D*_*BLOCK*_) and the variance among *D*_*BLOCK*_, hereafter Var[*D*_*BLOCK*_], to test for an excess in variance among genomic blocks that would be consistent with the presence of correlated genomic segments (haplotypes) with shared derived mutations. Under the null hypothesis of no introgression, Var[*D*_*BLOCK*_] among true genomic blocks derives largely from relatively small ancestral haplotypes such that random permutation of segments among blocks will not affect the variance among blocks. If the genome contains introgressed haplotypes that are larger than the segments, Var[*D*_*BLOCK*_] will be larger when these haplotypes are intact in the empirical data and smaller after random permutation that dissolves correlations among segments. We calculated Var[*D*_*BLOCK*_] among genomic blocks in the empirical data. Then, for each comparison, we permuted the internals ∽2.5 Kb segments among genomic blocks and re-calculated Var[*D*_*BLOCK*_] for each permuted genome. Then Var[*D*_*BLOCK*_] from the empirical data was compared to the distribution of Var[*D*_*BLOCK*_] from permuted genomes to ask whether variance among genomic blocks is higher in the empirical data where true correlations remain intact relative to the permuted genomes where variance in *D* will be driven largely by segregating ancestral haplotypes.

For the second test, establish confidence intervals for estimates of the *D*_*BLOCK*_ statistic in order to identify individual genomic blocks with significant evidence for introgression. To do so, we conducted block jackknife analyses within each genomic block [55] by dropping each genomic segment within a given block in turn and recalculating *D*_*BLOCK*_. We calculated 95% confidence intervals for each genomic block using variance estimated from this jackknife procedure. These confidence intervals are presented as ribbons in Figure 6.

For the third test, we established genome wide thresholds corrected for multiple testing in order identify genomic blocks exceeding these thresholds consistent with recent introgression. We conducted the permutation of segments within blocks procedure as above, but for each permuted genome, we calculated *D*_*BLOCK*_ and retained the maximum and minimum values of *D*_*BLOCK*_. To determine whether any individual true genomic blocks showed evidence of significant excess sharing of derived alleles, we established 95% critical thresholds (Table 2) from this permutation procedure and compared the value of *D*_*BLOCK*_ among true blocks. These genome-wide critical thresholds are presented as dashed lines in Figure 6.

To determine whether introgression has been very recent between *A. arabiensis* and either M form or GOUNDRY, we compared the proportion of the genome in windows with significant *D* values between sympatric *A. arabiensis* from Burkina Faso and allopatric *A. arabiensis* from Tanzania [38]. Since the standard assumption of introgression with only one of the two sister taxa holds for this test, we calculated the standard error of *D* for each comparison using the block jackknife approach and used a Z-test to assess significance [55,56].

### Comparing genetic divergence among genomic regions

To test hypotheses related to the role of recombination in determining the genomic architecture of reproductive isolation in this system, we divided the genome into regions based on expected levels of recombination in hypothetical hybrids. A fine-scale genetic map is not yet available for *Anopheles* mosquitoes, but it has been shown in *Drosophila* that recombination rates approach zero within several megabases on each side of the centromere and also near the telomeres [126,127]. Although patterns of linkage disequilibrium (LD) are also affected by processes other than local meiotic recombination rates, estimated recombination rate should give a rough approximation of expected LD across the genome. In fact, patterns of LD have been used to define genetic maps in some vertebrates and correspond approximately to genetic maps based on experimental crosses [128,129]. We measured background LD (see description of LD measurement above for details) in our M form and *A. arabiensis* samples, taking average *r*^2^ values within 10 kb physical windows across the genome. We found that LD was relatively constant across the genome except for large increases near the autosomal centromeres and smaller increases near the telomeres (Figure S2). Based on this pattern and the assumption that recombination rates in *Anopheles* correspond approximately to the *Drosophila* genetic map, we defined several broad recombinational categories for analysis. We first defined the ‘Pericentromeric-Telomeric’ regions of the autosomes to be all windows within 10 MB on either side of the centromere or within 1 MB from the telomere. It should be noted that we assumed that the starting and ending coordinates of the PEST reference chromosomal sequences were reliable indicators for distance from centromeres and telomeres. Unless a chromosomal inversion was present, all remaining regions on the autosomes were assigned to the ‘Freely Recombining’ category. For the comparison between *A. gambiae* and *A. merus,* we assigned all windows inside of the 2R*op* chromosomal inversion complex to the ‘Autosomal-Inversion’ category. We used the outer coordinates for 2R*o* and 2R*p* breakpoint regions estimated by Kamali et al. [40].

The X chromosome was categorized for each comparison, according to species-specific conditions. We did not define a general ‘Pericentromeric-Telomeric’ category for two reasons: 1) We did not observe an increase in LD in the euchromatic regions near centromeres and telomeres (Figure S2) similar to increases observed on the autosomes. 2) In *Drosophila* [126,127], the pericentromeric reduction in recombination affects a relatively small region on the X relative to the autosomes, and we have excluded a large heterochromatic region around the centromere that likely encompasses the effected region in *Anopheles.* For the comparison between *A. gambiae* and *A. merus*, and the comparison between the M and S molecular forms, the entire euchromatic region on the X was considered ‘Freely Recombining’ since no inversions differentiate these groups. For the comparison between *A. gambiae* and *A. arabiensis*, we assigned the entire euchromatic region of the X as ‘X-Inversion’, since these species are differentiated across nearly 75% of the entire chromosome and introgression rates have been estimated to be 0 in laboratory crosses [82]. For the comparison between GOUNDRY and the molecular forms of *A. gambiae*, the entire euchromatic X was categorized as ‘Freely-Recombining’ except for the region spanning 8.47 MB to 10.1 MB, which was categorized as ‘X-Inversion’. As described below, we were not able to identify inversion breakpoints for the GOUNDRY inversion, but these coordinates correspond to the outer boundaries of the region with reduced nucleotide diversity (Figure 4).

To identify regions that are barriers to introgression in the *Anopheles gambiae* species complex, we compared genetic divergence among the four genomic categories using the following logic. Since genome-wide variation in mutation rate and the effects of linked selection could also lead to genomic variation in divergence among species even in the absence of introgression, we jointly analyzed absolute genetic divergence (*D*_*xy*_) and nucleotide diversity (*π*) in 10 kb non-overlapping windows and asked whether differences in genetic divergence among genomic regions are observed that cannot be explained by differences in nucleotide diversity (where diversity approximates the effects of linked selection or variation in mutation rate). *D*_*xy*_, is an estimate of 2*μt* + 4*N*_*e*_*μ* where *N*_*e*_ is the effective size of the ancestral population, *μ* is the mutation rate, and *t* is the divergence time. *π* which provides an estimate of 4*N*_*e*_*μ* where *N*_*e*_ is the effective size of the current population. By jointly considering these, we can make comparisons among genomic regions that differ substantially in 4*N*_*e*_*μ* such that differences in *D*_*xy*_ largely reflect differences in 2*μt.* We note that this analysis assumes that estimates of 4*N*_*e*_*μ* from current populations (or in most cases, average of the estimates from the two subgroups) are reliable estimates of 4*N*_*e*_*μ* in the ancestral population, and we believe this to be a reasonable assumption given the recency of radiation of the *A. gambiae* species complex. Under scenarios with gene flow among diverging subgroups or species, *t* will be smaller in regions that have introgressed, so regional barriers to gene flow are expected to have especially large values of *t,* and therefore values of *D*_*xy*_ that are larger than regions with similar 4*N*_*e*_*μ* but more introgression. In contrast, under a model of divergence in allopatry with no introgression, *t* is approximately equal among genomic regions such that divergence should be determined largely by 4*N*_*e*_*μ*, and thus correlate well with nucleotide diversity, even if elevated. We made comparisons among genomic regions using this framework and assuming that most freely recombining autosomal regions will have been introgressed at some point in the history of divergence since high rates of recombination inhibit associations between barriers to introgression (e.g. hybrid sterility factors) and surrounding chromosomal regions. Therefore, we compared other genomic regions to freely recombining autosomal regions to with respect to both divergence and nucleotide diversity to ask whether these other regions harbor excess divergence consistent with less introgression in these regions. Comparisons of distributions of genetic divergence and nucleotide diversity were made using the Mann-Whitney test in R [112].

For this analysis, we used only 10 Kb windows at least 600 sites with data for the taxa involved. We chose to use a focal species comparison to minimize the number of comparisons for concise presentation, but we obtain similar results in other comparisons (GOUNDRY vs. *A. arabiensis,* for example) since most variation in divergence depends on variance in coalescence in the ancestral population. We note that estimates of nucleotide diversity in GOUNDRY are not reliable estimates of the ancestral population since GOUNDRY is partially inbred and has the large swept region on the X, so we used estimates of nucleotide diversity from the M form for this analysis.

### Estimating the age of the GOUNDRY X-linked selective sweep

We estimated the number of generations since the fixation of the haplotype inside the putative X*h* inversion region in GOUNDRY in the following way. We assumed that no new mutations increased to have a frequency > 50% after the selective sweep. Under this assumption, we can estimate the mean time since the most recent common ancestor of the sampled haplotypes, representing the time of the sweep, by assembling a consensus sequence among the haplotypes to represent the common ancestor and counting the number of mutations separating each haplotype from the ancestor. Then we can calculate the number of mutations divided by the total haplotype size and divide this number by the mutation rate to obtain the number of generations separating haplotype from the consensus. We summed the number of mutations across all diploid sequences, counting 1 for genotypes called heterozygotes and 2 for homozygotes, and divided by 2 times the total number of sites passing filters. This grand total was then divided by the mutation rate to get the number of mutations. This approach can be simplified and stated as the following where the age to the most recent common ancestor is

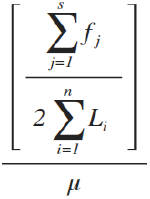

where *f*_*j*_ is the derived allele count at variable site *j*, *s* is the number of variable sites, *n* is the number of diploid individuals, *L*_*i*_ is the number of sites with called genotypes passing filters for individual *i*, and *μ* is the mutation rate, again assumed to be 7.0×10^−9^ (See demographic inference section above).

Genotypes for 12 diploid GOUNDRY individuals were queried across a region consisting of 1,372,538 sites after initial filtering. A total of 1,052 variable sites were recorded prior to additional read depth filtering. Since this estimate is highly sensitive to the number of mutations included, we estimated the number of generations using a series of filters varying in stringency. First, we removed clusters of variable sites, since these are likely to represent errors, by dividing the inversion region into 50 bp windows and excluding any windows with more than two variable sites (14 out of 33,478 windows). This resulted in a total of 989 variable sites across 1,367,970 sites in total. We also excluded sites with read depth in the top 5% for each individual according to each individual’s read depth distribution. Then we counted mutations for each individual using different thresholds such that sites passed filter if that individual was covered by at least 6, 8, 9, 10, 11, or 12 reads. We found that too few sites passed filtering to be informative with a 15-read cutoff. After conservatively filtering the data to minimize the effect of errors, the majority of the remaining variance in our estimates derives from variance in the number of mutations per haplotype. If we assume that the number of mutations per haplotype is Poisson distributed, we can calculate the standard deviation for read-filter point estimates by taking the square root of the point estimate, as the variance for a Poisson is equal to its mean.

Age estimates varied by 2.5 fold depending on the minimum read depth filter (Figure S12), decreasing from 1,975 generations to 776 generations with 6 and 12-read minimum cutoffs, respectively. However, excluding the 6 read cutoff, the remaining five age estimates varied by less than 1.4 fold, ranging from 776 to 1,079. Standard deviations largely overlap among estimates from the 8-12-read filters, suggesting that the estimates are quite similar in this range. If we examine how the depth filters affect mutation counts per site for each individual separately, the 6-read filter results in an increase in mutations for all individuals (Figure S12). The mutation counts per site remain relatively flat for the remaining filters with only a slight systematic downward trend with the 12-read filter. The estimated age drops substantially between the 6 and 8-read filters and continues to decrease slowly from there. This strongly suggests that the proportion of variable sites that are genotype-calling errors is relatively high for this region when low read-depth minimums are used. To determine whether genotype-calling errors may be inflating the estimated number of mutations, we calculated the minor allele count at each variable site for each of the read-depth thresholds and see that, when a minimum of 6 reads is used, there is an excess of doubletons compared to higher read depths (Figure S13), consistent with a high proportion of true heterozygous sites being called as homozygous for the minor allele. The ratio of doubletons to singletons decreases substantially when the minimum number of reads is increased to 8, implying that many erroneous homozygous-alternative genotype calls are converted to heterozygous with this threshold (Figure S13). This improvement does not fully explain the drop in the number of observed mutations, however.

We can leverage the extra sequencing invested in one individual to gauge the effect of our filtering process. Our sequencing effort was not evenly distributed among individual GOUNDRY samples since we sequenced one individual (GOUND-0446) to average 20.03x read depth, while the remaining individuals were sequenced to an average of 10.82x read depth. Since the accuracy of genotype calling is correlated with read depth, we expect the genotype calls made for this individual will harbor the fewest errors. When considering how the read-depth thresholds affect estimates from individual samples, GOUND-0446 showed patterns that differ from the other samples (Fig S12). Specifically, while the number of sites passing filter changes less for this individual with increasing read-depth minimums, the number of mutations observed in this individual is also less sensitive to minimum read-depth, suggesting that genotype calling is substantially more error prone in the lower-read depth individuals. The variance in the number of observed mutations among individuals reduces substantially when the 12-read minimum is implemented with GOUND-0446 falling in the middle of the distribution, despite variance in the number of sites passing filter remaining large. This suggests that many of the errors have been removed using this filter. We, therefore, convert the point estimate from this filter and the standard deviation from this estimate (see above) to years, conservatively assuming 10 generations per year [130], to obtain the estimated age.

### Permutation test of M-GOUNDRY divergence on the X

To specifically test whether the large sweep region on the X chromosome of GOUNDRY individuals is significantly more diverged than similarly sized windows on the X, we defined a new statistic to compare the average *D*_*xy*_ among 10 kb windows within the putative inverted region (n = 166) with the average *D*_*xy*_ among windows in every other 1.67 MB window on the X that doesn’t overlap with the inverted region (n = 1,499). The statistic *ΔD_xy_* is defined as

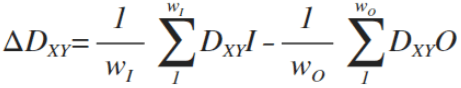

where *w*_*I*_ is the number of windows inside the inversion, *D*_*xy*_*I* is a *D*_*xy*_ value for a 10kb window inside the inversion, *w*_*O*_ is the number of windows outside the inversion, *D*_xy_*O* is a *D*_*XY*_ value for a 10kb window outside the inversion. A positive *ΔD_xy_* indicates that the average value inside the inversion is greater than the average value in the window outside the inversion. A negative value indicates that the average value is higher in the window outside the inversion. We calculated this statistic between M and GOUNDRY. For every window outside the swept region, we calculated *ΔD_xy_* with the true window assignments and then randomly permuted 10 Kb window assignments to the inversion or the outside window and recalculated *ΔD_xy_*. Each window comparison was permuted 10^5^ times.

One potential concern with this approach is that the windows inside of the inversion are part of a haplotype with little recombination while windows outside of the inversion represent many independent genealogical histories. However, this is a consideration only for very recent sequence evolution and most of the patterns of *D*_*xy*_ are determined by coalescence in the ancestral population prior to the origin of the inversion. After the inversion becomes relative common in the population, it will recombine normally with other inverted chromosomes in the population, so the comparison between windows inside and outside of the swept region should not be biased.

### Relative Node Depth expectation modeling

To explicitly test whether such a pattern could be obtained under a pure split model with no gene flow, we obtained expected values of Relative Node Depth (RND) by assuming a phylogeny where M form and GOUNDRY form a clade with S form as the outgroup and using coalescent theory under this model to calculate expected RND. We calculated the ratio of expected values of *D*_*GM*_ and *D*_*GS*_ as

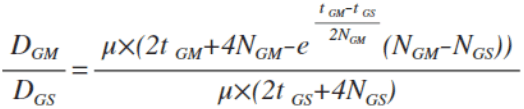

where *μ* is the mutation rate, *N*_*GM*_ and *N*_*GS*_ are the effective sizes of the M-GOUNDRY ancestral population and the M-S-GOUNDRY ancestral population, respectively, t_GM_ is the number of generations since the split between GOUNDRY and M form, and t_GS_ is the number of generations since the split between S form and the M-GOUNDRY. The denominator is the standard definition for *D*_*xy*_ [111], but we adjusted the equation in the numerator using standard coalescent theory to accommodate the probability that two the M and GOUNDRY lineages may either coalesce in the M-GOUNDRY ancestor or in the M-S-GOUNDRY ancestor that may differ in size, and therefore probability of coalescence, from the M-GOUNDRY ancestor (that may differ in size).

We calculated expected values of RND under a range of parameterizations using the M-GOUNDRY split time of 1,112,660 generations obtained from the demographic inference described above. For all calculations, we varied the M-GOUNDRY ancestral size from 100 to 10^6^, including 20,000 grid points. In the first set of calculations, we assumed that the split time between S and M-GOUNDRY was 1.1 times longer than the M-GOUNDRY split time and varied the relative sizes of the ancestral species such that the M-S-GOUNDRY ancestor such that the M-GOUNDRY ancestor varied from 1% to 100% the size of the M-S-GOUNDRY ancestor (Figure 10). We conducted a second set of calculations that were identical to the first except that we assumed that the difference in split times was 1.5 times longer instead of 1.1 (Figure 10). In the last set of calculations, we set the sizes of the two ancestors to be equal and varied the difference in split times such that the split time between S form and M-GOUNDRY was 1%, 10%, 20%, 40%, 70%, and 100% longer than the split time for M-GOUNDRY (Figure 10).

### Putative GOUNDRY Inversion breakpoint mapping

The dramatic reduction in nucleotide diversity across a 1.67MB region on the X chromosome in our GOUNDRY sample led us to hypothesize the existence of a novel chromosomal inversion in this region. Read mapping and depth is normal across the region containing the reduction of diversity (Figure S3). The exceptionally low level of nucleotide diversity in this region suggest a strong selective sweep recently fixed in this region, but such a wide footprint of the sweep and the remarkably rapid recovery to background diversity levels outside the sweep are both unexpected under normal recombination conditions. The existence of a new inversion that suppresses recombination with the alternative, presumably ancestral, form is the most likely scenario. We observe in our data the fixation of a 1.67 MB haplotype implying that this haplotype was maintained without recombination during the majority of the selective sweep. Since recombination should be normal among chromosomes carrying the inverted form, this suggests that the inverted form of the inversion was extremely rare in the population at the beginning of the sweep

Several issues inherent to this system and our data complicate identifying the breakpoints of this putative inversion. The first is that the inversion was recently under positive selection resulting in patterns of nucleotide diversity, linkage disequilibrium, homozygosity, and allele frequency differentiation that may extend beyond the breakpoints. There are clear changes in these population genetic signals in the putative region, but we can only conclude that the inversion lies somewhere inside of this footprint. The second issue is that GOUNDRY is inbred and harbors long tracts of IBD, some of which overlap the putative inverted region, making it difficult to distinguish homozygosity related to the sweep from that resulting from inbreeding. The third issue is that the X chromosome is repetitive and harbors many transposable elements resulting in a reference sequence riddled with gaps. Inversion breakpoints that have been characterized in *Anopheles* [131] often lie inside or near such lowly-complex, repetitive regions. Our short paired-end read data has insert sizes of approximately 470bps, so mapping the presumed breakpoints is expected to be challenging if they are in repetitive DNA, and we are unlikely to be able to map read pairs across these regions. It should be noted that very few chromosomal inversions identified with cytogenetics have been characterized molecularly, even in systems with reference sequences [40,103,132].

With these challenges in mind, we attempted to map the breakpoints of the inversion using a variety of approaches to collect candidate sites that could be assayed using PCR amplification. We manually inspected the short read data in this region using IGV [133] and located the edges of the swept region at positions 8,462,788 on the left and 10,137,178 on the right, according to coordinates in the AgamP3 PEST reference sequence. Since the selective sweep likely purged nucleotide diversity from the region surrounding the inverted haplotype as well, we attempted to identify the true breakpoints assuming that true breakpoints fall just inside these boundaries. For all of the following analyses except the read-depth analysis, we used reads that were trimmed using SolexaQA (v.2.1; [134]) with default settings and 50bp minimum size. We attempted a large series of *de novo* assemblies using the multi-kmer approach in SOAPdenovo2 [135] with –R –F and –p 4. We first attempted a de novo assembly of just reads that mapped to the inversion region on the X and obtained many contig sets based on variety of subsets of the data. None of the contigs showed the ‘T’ shape expected when the contig is aligned to the PEST reference and part of the contig is inverted. We also looked for cases of consensus where a variety of de novo assemblies using subsets of the data or all GOUNDRY reads all failed to assemble across a point on the reference, assuming that we could rule out regions that were properly assembled and aligned and focus on regions that fail to assemble as candidates. In another approach, we measured mapped read depth in 100bp windows across the region in GOUNDRY, M form, and *A. arabiensis*. We then systematically searched for windows where M form and *A. arabiensis* showed normal read depths while GOUNDRY reads failed to map in any sample. We also took a similar mapping-based approach where we identified all reads where the pair was mapped in the incorrect orientation or only one of the two mates mapped. We then searched for local enrichments of these mate-pair violations. Lastly, we used two computational implementations of algorithms specifically designed to identify structural variants such as inversions. We used PINDEL [136] with a window size of 2 million and a minimum inversion size of 2000. We also used GASV [132] with default settings.

From each of these approaches, we identified a series of candidate breakpoints. Several of the candidates were supported by multiple approaches. For these candidates, we designed primers on either side and attempted to PCR amplify across the candidate region in samples of GOUNDRY, M form, and *A. arabiensis*, with the assumption that a true breakpoint should amplify in M form and *A. arabiensis*, but not in GOUNDRY. However, all of the candidate PCR reactions either amplified in all three groups or failed to reliably amplify in any group, ultimately preventing the molecular characterization of this inversion.

## Acknowledgements

We thank Matteo Fumagalli, Filipe Vieira, and Tyler Linderoth for assistance with next generation sequence data analyses and ANGSD. We thank members of the Nielsen group for helpful discussions on various aspects of this work. We also thank Russ Corbett-Detig and three anonymous reviewers for helpful comments on an earlier version of this manuscript. We are thankful for the use of the Extreme Science and Engineering Discovery Environment (XSEDE), which is supported by National Science Foundation grant number OCI-1053575.

## Supplementary Table Legends

**Table S1**: Next-generation sequencing statistics for mosquito samples.

**Table S2**: Inferred selective sweeps in GOUNDRY, M form and *A. arabiensis*. Chr: chromosomal location of sweep; Peak Pos: location of window with highest LLR; LLR – log likelihood ratio from SweepFinder; *P*_gen_ – genome-wide *P* value, 0 values indicate *P* values less than 0.005; *P*_*loc*_ – per-locus *P* value, 0 values correspond to *P* values less than 1.26×10-6 for autosomes and less than 6.67×10-6 for the X chromosome. Gene name, ImmunoDB ID, and Gene description from Vectorbase.org.

**Table S3:** Genes located inside large swept region (∽8.47 – 10.1 MB) on X chromosome in GOUNDRY subgroup of *A. gambiae*. Chr: chromosomal location of sweep; Gene name, ImmunoDB ID, and Gene description from Vectorbase.org.

**Table S4:** Genes located inside introgressed windows with genome-wide significant values of *D.* Introg - Introgression between indicated subgroups; G_Abf – Introgression between GOUNDRY and Burkina Faso population of *A. arabiensis*; M_Abf - Introgression between M form and Burkina Faso population of *A. arabiensis*; M_Sform - Introgression between M and S forms; G_Sform - Introgression between GOUNDRY and S form; Chr – chromosomal location; Gene start bp – basepair coordinate of start in coding gene sequence; Gene end bp – basepair coordinate of end in coding gene sequence; Gene name, ImmunoDB ID, and Gene description from Vectorbase.org.

**Table S5:** Collection site and date information for mosquito samples.

**Table S6**: Summary statistics for mapping to *A. gambiae* PEST reference and subgroup SPEC reference.

**Table S7**: Sites included and excluded from analysis in all subgroups.

## Supplementary Figure Legends

**Figure S1:** Test for differences in the amount of recent selection between the GOUNDRY versus M form subgroups. We tested whether the number of megabases harboring at least a single significant selective sweep window differed between the subgroups by subtracting the number of megabases in M form from the number of megabases in GOUNDRY. The red line indicates the true difference. The histogram presents the same calculation made using 10^4^ datasets with subgroup assignments randomly permuted. No permuted genome produced the same or larger difference between the two subgroups.

**Figure S2:** Background Linkage Disequilibrium (LD) for each chromosomal arm in *A. gambiae* M form and *A. arabiensis*. LD (*r^2^*) between SNPs separated by at least 1kb (10 kb maximum) was averaged in 10 kb non-overlapping windows and LOESS-smoothed using a span of 1%. Low complexity and heterochromatic regions have been excluded. The large spikes on 2L and X (noted with vertical blue shaded bars) also coincide with reductions in nucleotide diversity and are thus likely to be recent selective sweeps. Additionally, there are a number of long distance increases (noted with horizontal grey bars) in LD in both M form and *A. arabiensis* that coincide approximately with the locations of known chromosomal inversions segregating in these subgroups [41].

**Figure S3:** Mean total read depth for GOUNDRY X chromosome sweep region. Mean total read depth across all GOUNDRY samples (n=12) for sites within non-overlapping 500 bp windows and plotted as a function of chromosomal position (megabases). The position of large GOUNDRY X sweep region is shown with grey bar.

**Figure S4:** Relative genetic divergence (*D*_*a*_) between GOUNDRY and M form. *Da* plotted as a function of M form nucleotide diversity using only intergenic sites in non-overlapping 10kb windows. Low complexity and heterochromatic regions were excluded. Genomic regions were defined based on predicted rates of recombination in hybrids (see Methods) and compared using non-parametric Mann-Whitney tests. X-Free: freely recombining regions on X chromosome. X-Inv: region inside putative X*h* chromosomal inversion. Asterisks indicate P values less than 4.896e-08.

**Figure S5:** *ΔD_xy_* between *A. gambiae* subgroups GOUNDRY and M form plotted for windows across the X. Permutation test *P*-values for GOUNDRY vs. M form comparisons presented on log-scale in bottom panel with standard and Bonferonni-corrected thresholds (Methods). Grey bar indicates inverted X*h* chromosomal region.

**Figure S6:** Autosomal 1D unfolded site frequency spectra (SFS) for intergenic variable sites in each subgroup. The SFS expected under standard coalescent equilibrium conditions is presented in black. The SFS was inferred for M form and *A. arabiensis* using –realSFS 1 (see Methods), but the SFS for GOUNDRY was inferred using the inbreeding-aware version (-realSFS 2) after estimating inbreeding coefficients for this subgroup (see Methods).

**Figure S7:** Inbreeding coefficients for each individual and each chromosomal arm. Inbreeding coefficients were estimated directly from genotype likelihoods for each chromosomal arm separately using Vieira et al. 2013 (see Methods). Each panel corresponds to a chromosomal arm. Each bar represents an individual mosquito sample. Colors indicate subgroup assignment according to the legend.

**Figure S8:** Decay of Linkage Disequilibrium (LD) in *A. gambiae* M form and *A. arabiensis*. LD (*r^2^*) between SNPs separated by no more than 5 kb binned, averaged, and plotted as a function of physical distance. Low complexity regions were excluded. Chromosomal arms 2R, 3L, and 3R were included for the autosome curves and the X plotted separately. Note different X and autosome y-axis scales

**Figure S9:** Comparison between the data, the best-fit model from *dadi*, and neutral coalescent simulations using MaCS. Site-frequency spectra were calculated from the **A)** true autosomal data, **B)** best-fit autosomal demographic model inferred using *dadi*, and **C)** polymorphism data generated using neutral coalescent simulations in MaCS. LD decay curves from true data (red line) are compared to decay curves inferred from **D)** coalescent simulations for X chromosome (grey), and **E)** coalescent simulations for autosomal data.

**Figure S10:** A) Distribution of maximum SweepFinder log-likelihood ratio statistics in a 50 kb region of neutral polymorphism data simulated under the demographic model inferred for GOUNDRY (top) and M form (bottom). See Methods for simulation details. B) Distribution of maximum SweepFinder log-likelihood ratio statistics in a full synthetic genome of neutral polymorphism data simulated under the demographic model inferred for GOUNDRY (top) and M form (bottom). See Methods for simulation details.

**Figure S11:** A) A representative cluster of significant selection peaks from Sweepfinder in GOUNDRY that were collapsed. The top panel shows the LLR profile when SweepFinder was applied to only variable sites (Red = genome wide significance; Gold = per-locus significant; Grey = not significant). The bottom panel presents the LLR profile when SweepFinder was applied to all sites. Critical thresholds were not established for ‘all-site’ analyses. The different y-axis values reflect the different sets of data considered in the composite likelihood function. The dotted line on the bottom panel indicates our threshold for ‘low’ values (LLR = 20). Under our clustering approach, this cluster was collapsed into a single peak. B) A representative cluster of significant selection peaks from Sweepfinder in GOUNDRY that were not collapsed. The top panel shows the LLR profile when SweepFinder was applied to only variable sites (Red = genome wide significance; Gold = per-locus significant; Grey = not significant). The bottom panel presents the LLR profile when SweepFinder was applied to all sites. Critical thresholds were not established for ‘all-site’ analyses. The dotted line on the bottom panel indicates our threshold for ‘low’ values (LLR = 20). The different y-axis values reflect the different sets of data considered in the composite likelihood function. Under our clustering approach, this cluster was not collapsed into a single peak.

**Figure S12:** Estimating the age of the selective sweep inside the putative (X*h*) inversion on the X chromosome in GOUNDRY. **A)** Each line shows the number of sites in millions that pass filtering according to different minimum read depth filters. **B)** The number of mutations counted for each individual according to minimum read filters. **C)** The average age in generations of the two chromosomes in each diploid individual as function of minimum read depth. **D)** Estimates for age of the most recent common ancestor as a function of minimum read depth. Bars indicate standard deviations as calculated from the average number mutations per haplotype. Colors correspond to individuals and are the same among panels A-C, with GOUND-0446 indicated in light-blue.

**Figure S13:** Minor allele counts at variable sites within the large X-linked sweep region (X*h*) in GOUNDRY genomes. Counts were made using called genotypes. Each panel presents a histogram distribution of allele counts at sites covered by at least 6-12 reads per individual (see Methods).

